# Birth synchronisation increases allonursing opportunities in free-ranging dogs

**DOI:** 10.64898/2026.10.11.758287

**Authors:** Melissa Vanderheyden, Nina Truffaz, Brenda Chaignon, Clément Car, Małgorzata Pilot, Giulia Cimarelli, Sarah Marshall-Pescini, Friederike Range, Andreas Berghänel

## Abstract

Over 50 years ago, the hypothesis emerged that co-resident women synchronise menstruation through pheromonal cues. This idea has persisted as a widespread narrative and has subsequently been extrapolated to other animals as well. Indeed, reproductive synchrony has many potential benefits, including creating allonursing opportunities and facilitating female mate choice. Nevertheless, little evidence exists of female vertebrates synchronising their oestrus beyond seasonal effects, through socially mediated cues. We analysed reproductive synchrony in free-ranging dogs, which combine weak, population-dependent reproductive seasonality with facultative cooperative breeding. Controlling for seasonality, we show that birth synchrony was higher among co-resident, maternally related females, suggesting synchronisation among individuals who are likely to cooperate. This was supported by our finding that birth synchrony and maternal kinship independently increased the probability and frequency of allonursing. Finally, preliminary evidence shows that oestrus synchrony decreased male monopolisation. These findings contribute to the debate on socially mediated reproductive synchronisation, suggesting that it may have mainly evolved because it facilitates cooperative breeding.

## 1. Introduction

After McClintock [1] reported in 1971 that human female friends align their menstrual cycles, the idea of oestrous synchronisation has provoked both popular myths and scientific debate [2–4]. Despite methodological criticism and a lack of conclusive support, the belief in human menstrual synchronisation persists demystification [5]. Still, convincing evidence of socially mediated oestrous synchrony in animals, particularly vertebrates, is scarce.

This paucity of evidence may be due to the difficulty to differentiate socially mediated synchronisation from seasonal reproduction following environmental cues. Seasonal breeding is common among animals, allowing reproduction to align with favourable environmental conditions [6,7]. Small seasonal time windows relative to a female’s receptive period lead to high synchrony [8], whilst a wider time frame promotes asynchrony [9]. For a wide range of animal species, however, reproductive synchrony delivers benefits that extend beyond adaptations to seasonality. Synchronous mating, for example, can prevent a single dominant male from monopolising fertile females, as described by the priority of access model [10]. Empirical studies on primates demonstrate that being able to mate with multiple males allows females to decrease the mating skew [11] and create paternity confusion, thereby removing the incentives for infanticide [12–14]. Additionally, females are able to express mate preferences [15] and are more likely to conceive [16].

Synchronous births, on the other hand, may be adaptive by facilitating predator swamping [17], joint protection of infants [18] and other forms of allomaternal care, in particular allonursing [19]. Hence, birth synchrony may facilitate cooperative breeding, here broadly defined as ‘a reproductive system in which helpers – i.e., individuals other than the parents – cooperate to raise offspring and thereby increase the immatures’ growth and survival ([20], p. 1; see also [21]). By facilitating cooperative care, reproductive synchrony may unlock new evolutionary pathways. In some eusocial ants, labour division is thought to be preceded by synchronised reproductive cycles [22], and cooperative breeding is thought to create favourable conditions for the evolution of socio-cognitive skills in primates including humans [23].

Since many benefits of reproductive synchrony are independent of seasonal effects, one might expect natural selection to lead to numerous cases where synchrony exceeds what is dictated by the seasonal time window, and is socially mediated through cues between females rather than solely by environmental cues [24]. In mammals, synchronising cues between females have been suggested to be olfactory [25,26], resembling the oestrous- or abortion-inducing olfactory cues emitted by male mice (Bruce and Whitten effect) [27]. Olfactory cue transmission is likely higher between individuals that interact more frequently. Socially induced reproductive synchrony between individuals that associate regularly was initially reported in humans [1] and subsequently in rodents [28,29], ruminants [26], carnivores [30], and primates [31]. However, after methodological criticism [2–4,30,32], these results could seldomly be replicated (e.g. [33–36]). In addition to seasonality [37], apparent synchronisation may arise from statistical artifacts [2–4,30,32], reproductive reset after synchronous offspring loss [30] or coordinated mating [38], rather than from physiological entrainment of the oestrus cycle.

Unambiguous evidence for socially mediated birth synchronisation is currently limited to two rather exceptional cases: the banded mongoose (*Mungos mungo*) and greater ani (*Crotophaga major*) [39– 41]. Both are obligate cooperative breeders; parents cannot successfully raise offspring without helpers. In contrast to the typical obligate cooperative breeding system with a dominant breeding pair and reproductive suppression of subordinate helpers, cooperation in banded mongooses and greater anis occurs between multiple breeding females. Females share a den or nest and give birth largely on the same day. Individuals trying to gain a competitive advantage for their offspring by giving birth ahead of others are punished through infanticide [39–41]. Hence, extreme birth synchronisation in these species is a prerequisite for maintaining the cooperative system by preventing single individuals from cheating and exploiting it. Since obligate cooperation between multiple breeding females is exceptional, it remains an open question whether synchronisation can also occur in breeding systems where selective pressures are not as high, e.g. where cooperative breeding is facultative.

Canid social systems present enticing subjects for investigating this question. They range from single breeding pairs providing biparental care, with or without help from non-breeding group members, to groups containing multiple breeding females [42–44]. Within-species flexibility is high, and whilst breeding in pairs is most common among canids, additional females may breed when ecological conditions allow for it. Multiple breeding females may allonurse and provide allomaternal care together, and they regularly receive help from males or non-breeding individuals [42–44]. To facilitate alloparental care, many canid species exhibit adaptations such as allofeeding through regurgitation and pseudopregnancy, which hormonally stimulates maternal behaviour and lactation [43,45]. The main challenge to study socially mediated synchronisation in canids is to differentiate the effects of synchronisation from their strict reproductive seasonality, which may even encompass a spermatogenesis pause outside of the breeding season [25,42,43,46]. Here, we tackle this challenge by leveraging the exceptional characteristics of free-ranging mongrel dogs.

Free-ranging dogs can live in multimale-multifemale groups of up to 27 individuals with multiple breeding females [47–50]. Unlike other canids, they are polygynandrous and reproduce twice a year [42,50]. Seasonality is absent in some populations and weak in others, depending on the local climate [42,47,48,51]. Alloparental care is predominantly provided by the mother’s female relatives and includes communal denning [52], allonursing, food offering, food regurgitation, protection, grooming, play and thermoregulation through body contact [49,53]. Without social monogamy, paternity is uncertain and males rarely provide care, except for protection and play [49,53]. For allonursing in particular, a minimum level of synchrony is required, given the comparatively short lactation period of max. 7 to 10 weeks [53]. Synchrony may be caused by seasonality in some populations [51], but even in populations without clear seasonality such as ours, overlapping pregnancies within social groups and allonursing are common [54]. Nevertheless, birth synchronisation has never been investigated in this species.

We studied reproductive synchronisation in a large population of free-ranging mongrel dogs in Morocco, investigating whether birth synchrony increases with increasing kinship and social relationship strength whilst controlling for potential seasonal effects. We did so first on the population level, i.e., across groups, and second among group members only, to address within-group patterns in greater detail. Subsequently, we tested whether synchrony and maternal kinship increase the likelihood that allomaternal care occurs within a dyad, and additionally whether higher synchrony also leads to higher frequencies of allomaternal care. Finally, we investigated whether increased oestrous synchrony relates to reduced monopolisation by single males, reflected in an increasing number of mates per female.

Based on the hypothesis that socially mediated synchrony is adaptive in dogs by facilitating allomaternal care, we expected to find higher birth synchrony among females who are more likely to cooperate: members of the same social group who are related [19,55–57] or have a strong social relationship [58]. Additionally, we expected more synchronous dyads from the same group to be more likely to care for each other’s offspring. Alternatively, if females synchronise to avoid monopolisation during oestrous, we expected them to mate with more males when a group member is in heat simultaneously.

## 2. Methods

### (a) Study population and data collection

Observational data were collected through scan and focal observations in our study area in Tamraght, a coastal town in the Souss-Massa region, Morocco (30°31′N, 9°41′W). The individuals belonged to a stable population, monitored since 2016 (e.g. see [59]). The climate is mild, with limited contrasts between winter and summer, and monthly temperatures reaching average maxima of 20°C to 29°C and average minima of 11°C to 20°C. Mean monthly precipitation from October to March is 19 mm, with near-zero rainfall during summer months. The dogs in the study area scavenge on human-generated food sources. Most are habituated to people, as tourists and locals feed and interact with them. Genetically, the population clusters closer to other free-ranging dog populations than to purebreds [60], indicating that it derived from a continuous lineage of free-breeding dogs rather than from an admixture of breeds, similar to Eurasian free-ranging dog populations [61].

Proximity scan observations [62] were carried out along transects of predetermined areas (methods described in [50]). These yielded GPS locations for individual dogs, which were identified based on individual morphology (Supplementary Fig. S1). Individuals outside the scan area were also recorded if located within 20 meters of a dog inside the scan zone. The order of scan areas was randomized across transect walks, to reduce spatial and temporal dependence. GPS locations were recorded using the Map Marker app on a handheld tablet and used to calculate dyadic distances between individuals. During the period included in this study, from 22 April 2022 to 31 December 2024, scan observations usually took place once daily, adding up to a total of 1102.

Focal data [62] were collected for 1129 hours in total, on 38 adult females which were part of eight stable social groups between 15 March 2023 and 28 March 2024 (29.7 ± 14.3 hours per individual). We focused observation efforts on the hours after sunrise and before sunset, since dogs were less active during the midday heat. Each focal observation lasted 15 minutes, during which we documented all occurrences of social interactions involving the focal animal, including proximity of less than 2 body lengths and affiliative and agonistic behaviours. Allomaternal care was either recorded as part of a focal observation or collected ad libitum during either the scan or the focal observations. Behaviours defined as care were nursing, grooming, lying in body contact and play, though play was never observed in this context. The rare event of pups attempting to suckle from non-lactating females was recorded as ‘body contact’.

Between 15 November 2023 and 18 March 2024, we additionally carried out 200 hours of focal observations of 8 females in heat, recording all occurrences of male-female interactions. We selected these females from the individuals included in the basic focal observations described above, based on visual signs of heat (swollen vulva, discharge, attraction of males). When two females from the same group were in heat on the same day (on 8 of 40 observation days), we followed the first female to enter heat. If the second female stayed in heat after the first female had finished, we switched to her.

### (b)Birth synchrony calculation

Birth dates were recorded ad libitum (raw data example in Fig. 1A). We included litters born between 22 September 2022 to 12 May 2025, for which scan or focal data on the mother were available. We recorded the size and GPS location of each litter, assessing the birth date based on the body condition of the pups. Parturition dates with an estimation error larger than ± 7 days were excluded. To calculate reproductive synchrony, we constructed dyadic combinations of all included litters (details in Supplementary Text, Supplementary Fig. S2). Reproductive synchrony was defined as the absolute difference in days between the parturition dates of two mothers, with larger values indicating lower synchrony.

**Fig 1.**
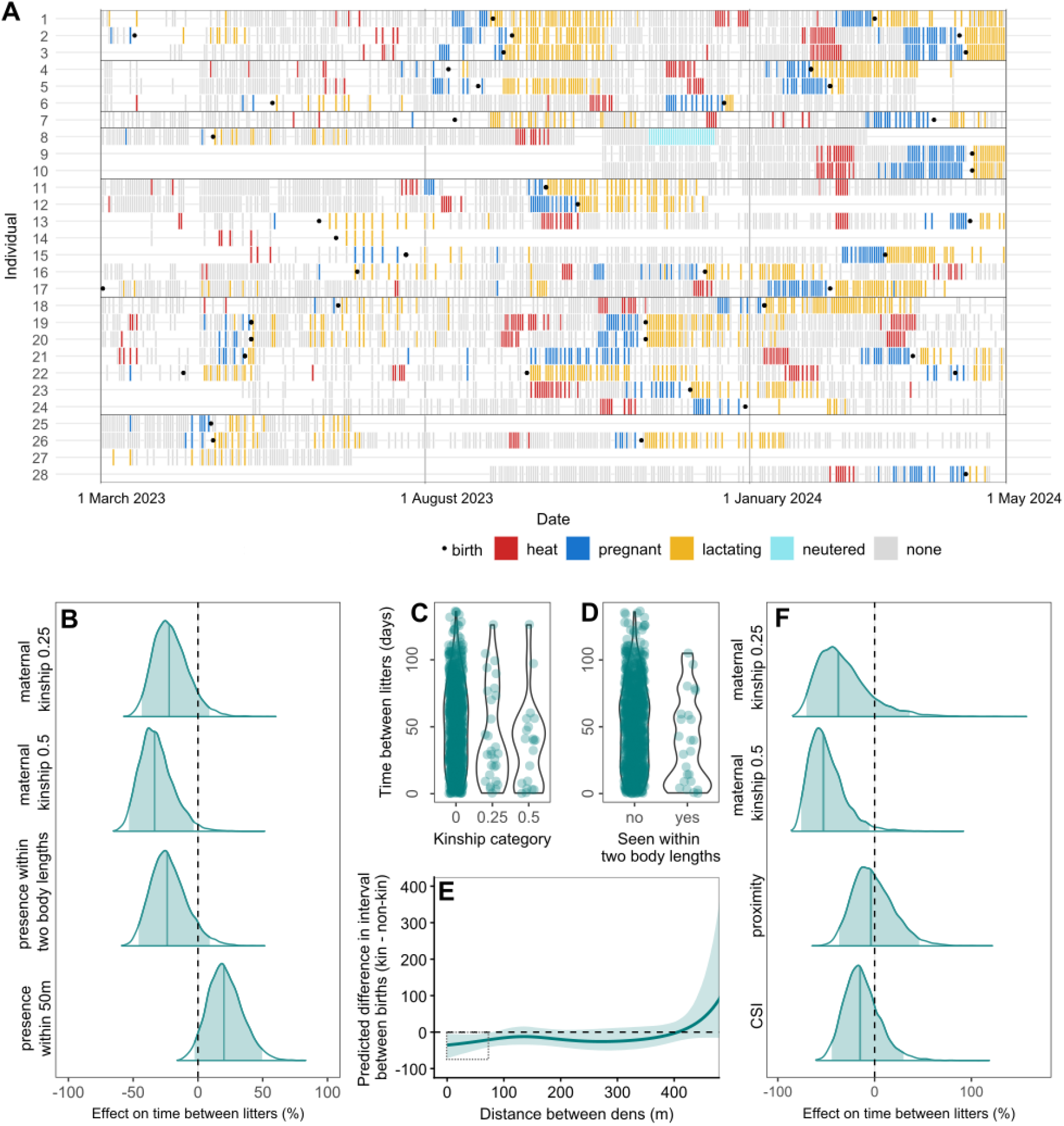
Reproductive synchrony at the population level and within groups. (**A**) Visualisation of raw reproductive data per individual. Rectangles are days on which the individual was observed. Full circles represent litter births. Neutering is shown for multiple days after the procedure to enhance visibility. Large boxes delineate social groups. Only females with at least one reproductive state or event are included. Only focal females during the focal observation period, for which the most complete dataset was collected, are shown. For a visualisation with all females included in this study see Supplementary Fig. S10. (**B - E**) Synchrony among dyads across the population. (**B**) The effect of relationship strength and kinship on the time between births, using scan proximities (within two body lengths and within 50 m) to approximate relationship strength (N = 898 litter combinations; posterior distributions from Bayesian Gamma mixed-effects models). Reference category for maternal kinship: non-kin or unknown. Solid lines: posterior means. Shaded areas: 95% credible intervals. Dashed line at zero: no effect. (**C, D**) Raw data violin plots showing the time between litters as a function of maternal kinship and presence within two body lengths. Each point represents a litter dyad. (**E**) The effect of maternal kinship on synchrony decreases with increasing distance. Posterior predictions from the Bayesian Gamma mixed-effects model with separate smooth functions for kin and non-kin dyads. The solid line is the posterior mean difference between the kin and non-kin groups (kin minus non-kin), which shows that synchrony was higher for kin dyads, but only at short distances (<±100 m, indicated by the grey box). Shaded areas indicate 95% credible intervals. For the illustration, maternal kinship was held constant at zero and random effects were excluded. (**F**) Synchrony among dyads from the same group. Posterior distributions from the Bayesian Gamma mixed-effects model investigating the effect of relationship strength and maternal kinship (0.25 and 0.5, in comparison with non-kin) on the time between births (N = 35 litter combinations). Proximity: time spent within two body lengths, corrected for the number of observations as described in Methods. CSI: Composite Sociality Index. Solid lines: posterior means. Shaded areas: 95% credible intervals. The dashed line at zero denotes no effect.

### (c) Kinship from demographic and genetic data

We used two kinds of kinship data: maternal kinship derived from demographic observational data and genetic kinship obtained through DNA analysis. This distinction reflects the possibility that dogs recognise maternal but not paternal kin, given the polygynandrous mating system with extra-pack paternity and high rates of multiple paternity [50]. Moreover, as demographic kinship could be obtained for more dyads than genetic kinship, we prioritized investigating the effects of maternal kinship. Subsequently, we used the available genetic kinship data to further analyse the effect of overall relatedness. Maternal kinship was included as a factor with three categories of expected relatedness: 0.5 (mother-daughter), 0.25 (e.g. grandmother-granddaughter) and 0 (non-kin or unknown), whilst genetic kinship was a continuous variable (details in Supplementary Text).

### (d) Social relationship strength calculation

We calculated dyadic social relationship strength between mothers using two distinct data sets. First, we investigated synchrony across the general population, using scan data, then at a finer scale within social groups, using focal data.

With the scan data, we approximated the relationship strength between two mothers as their (binary) presence within 50 meters and within two body lengths of each other (details in Supplementary Text). These distances were assumed to identify females who were group members or close neighbours, and those with a close social relationship, respectively. Using the focal data, dyadic relationship strength was calculated based on 1) the frequency and duration of affiliative interactions (body contact, grooming, play, greeting and friendly postures, combined into the composite sociality index, CSI) and 2) the time spent within two body lengths (details in Supplementary Text). We distinguished between affiliative interactions and close proximity since they potentially represent different dimensions of sociality (e.g. proximity can result from two individuals being attracted to a third individual or to a food source) [63]. Hence, the CSI without proximity may better distinguish strong from weak relationships.

### (d) Statistical analysis

All statistical analyses were done with Bayesian generalised mixed-effects models, using the ‘brms’ package in R [64,65]. Model convergence was assessed using 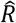 (≤1.01), effective sample sizes and Hamiltonian Monte Carlo diagnostics. Model fit was evaluated using posterior predictive checks. For the synchrony and allomaternal care analyses, random-effect factors included the mother dyad, as well as the identities of the two mothers and the two litters involved. Because individuals could appear as either member of a dyad, the random effects for mother and litter identity were specified as multi-membership effects, allowing each observation to be associated with both individuals (or litters) regardless of their order within the dyad. For the male monopolisation analysis, the random effects factor was focal female identity. All continuous predictors were z-transformed. Weakly informative priors were defined based on previous knowledge about the expected direction of effects, with wide uncertainty encompassing zero as well as large effects in the proposed direction. The intercept prior was based on the average of the response variable ± 2 SDs. Collinearity among fixed predictors and its consequences on model accuracy was assessed using generalised variance inflation factors (GVIF) calculated from corresponding linear models, using the package ‘car’ [66]. All GVIFs were below 1.5, indicating negligible consequences on estimation precision.

#### Model 1 and 2: Effect of kinship and social relationship on synchrony

Two models were run to test the influence of kinship and social relationship strength on reproductive synchrony. In Model 1, social relationship strength was based on proximity data from scan observations. In Model 2, social relationship strength was based on affiliative behaviours and proximities recorded during focal sessions. Since synchrony was measured as the absolute number of days between two births, it was cut at zero. Therefore, both models were fitted with a Gamma distribution and logarithmic link function, with synchrony values of exactly zero days entered as 0.5 days. Seasonal patterns were controlled for by including a non-linear cyclic effect of day number within the year (1 to 365), modelled with a cubic spline function allowing for high wiggliness (k = 10) to control for potential seasonal effects thoroughly even if they follow a complex pattern. We used a Normal (4.2, 2) prior on the log scale for the intercept and a Normal (-0.5, 0.5) prior for all predictor variables.

For Model 1, the predictor variables were successively added in 3 steps because a) we wanted to run a mediation analysis, investigating whether potential effects of social relationship strength were mediated by kinship (relatives may prefer each other as social partners [67]) and b) genetic kinship was known for only 52% of the original sample size. First, we ran a basic model including only the two measures of relationship strength to investigate their isolated effects (N = 898 litter combinations with 108 litters from 64 females). Second, we added maternal kinship as categorical variable (0, 0.25 and 0.5) to investigate its effect and whether it mediates potential effects of relationship strength (N = 898 litter combinations). Third, we further added genetic kinship as a continuous variable. Due to the large number of dyads for which genetic data were unavailable, this reduced the sample size by half (N = 432 litter combinations with 75 litters from 39 females).

We additionally tested how the distance between dens influenced synchrony depending on maternal kinship status by modelling the relationship between synchrony and den distance using group-specific smooth functions in a Bayesian Gamma mixed-effects model. Group-specific smooth terms were used to allow for nonlinear effects of distance and were estimated separately for dyads with and without maternal kinship (binary). We used cubic spline functions with constrained wiggliness (k = 4) and defined moderate priors for the standard deviation (class “sds”) and thus the wiggliness of the smoothing splines using a t-distribution with student_t (1, 0, 2). Only dyads with dens within 500 meters were included; since kin dyads are rare (0.06%) at larger distances, comparisons with non-kin are only meaningful within this range (N = 180 litter combinations with 85 litters from 54 mothers). To identify the den distances at which the two group-specific smooth functions diverged, we estimated the posterior smooth functions for each group using posterior_epred in brms and subsequently calculated and plotted the difference between them together with its corresponding credible interval.

Model 2 on focal observation data included only dyads of focal females belonging to the same group (N = 35 litter combinations with 31 litters from 23 females). Genetic kinship was only known for 8 out of the 30 included mother dyads, and therefore not used as a predictor variable.

#### Model 3 and 4: Effect of synchrony and kinship on allomaternal care

We tested whether, within social groups, dyadic synchrony and kinship increased the probability of allomaternal care being provided by either mother in the dyad to at least one pup during its first three months of life. We analysed the effects of synchrony and kinship on nursing separate from those on other forms of allomaternal care (allogrooming and lying in body contact). The outcome variables were undirected, not distinguishing which of the two mothers provided care. Dyads were included if at least one pup from each litter survived longer than two weeks. Only maternal kinship was included as a predictor variable, since genetic kinship was unknown for most dyads.

Higher synchrony may promote the occurrence of allonursing and allomaternal care but not necessarily increase their frequency. Therefore, Model 3 investigated the effects of kinship and synchrony on whether allomaternal care or allonursing occurred at all. Using ad libitum records of allomaternal care collected during both focal and scan observations allowed us to explore the general occurrence of allomaternal care in the largest dataset possible (N = 33 litter combinations with 42 litters from 32 females). The outcome variable was the presence or absence of allomaternal care per dyad, and the model was fit with a Bernoulli distribution and a logit link function. The data source (focal or scan observations) was included as a control variable. For the intercept, we used a Normal (0, 2) prior on the log odds scale. Predictor and control variables for which a negative effect was expected (‘time between births’ and ‘source’) were assigned a Normal (-2, 2) prior, whilst we used a Normal (2, 2) prior for variables with an expected positive effect (maternal kinship of 0.5 and 0.25).

Model 4 additionally investigated the effects of kinship and synchrony on the frequency of allomaternal care. Frequency was calculated as the proportion of 15-minute focal sessions in which allomaternal care occurred, limiting the sample size to focal females (N = 23 litter combinations with 22 litters from 17 females; full methodological details in Supplementary Text).

#### Model 5: Effect of synchrony on the number of mating partners

We studied how oestrus synchrony impacts male monopolisation potential by testing the effect of the number of fertile females in a social group on the number of males that mated with the focal female on a given day, excluding days where no mating was observed (N = 26 days, 8 females). A Poisson model with log link was used.

## 3. Results

### (a) Population-level birth synchrony is higher among maternal kin

Across the population (Model 1), we found significant seasonal variation in birth synchrony, with the cyclic smooth term showing non-zero wiggliness (sds = 0.23, 95% CI 0.10–0.48, Supplementary Fig. S3). However, births occurred throughout the year, and visual inspection shows only weak seasonality (Supplementary Fig. S4). Comparing the model before and after adding maternal kinship as a predictor shows that maternal kinship partially mediated the effect of social relationship strength, i.e. the synchronising effect of being observed in close proximity (two body lengths) was partly explained by maternal kinship and weakened after including the latter (reduced model results in Supplementary Fig. S5 and Table S1; full model results and raw data in Fig. 1 B-D, Supplementary Tables S2A and S3). Yet, both predictor variables had a synchronising effect in the full model.

In detail, the time between births was 22.1% shorter for mother dyads with maternal kinship of 0.25 (95% CI: −43.4 to 8.3%; pd: 93.1%), and 33.6% shorter for dyads with maternal kinship of 0.5 (95% CI: −53.2 to −3.0%; pd: 98.4%), compared with dyads that were unrelated or for which maternal kinship was unknown. Mothers gave birth 23.7% closer in time when they were observed within two body lengths of each other compared with mothers that were not (95% CI: −46.1 to 9.4%, pd: 93.3%). Given the results of the mediation analysis, this remaining association between close proximity and birth synchrony after controlling for maternal kinship may be caused by dyads that were maternally related but for which kinship could not be identified and hence were included as ‘unrelated’. In these cases, close proximity could act as a proxy for unobserved relatedness rather than representing an independent effect. Against our predictions, mother dyads that were observed within 50 meters of each other had 20.9% longer gaps between births than dyads that were not seen together (95% CI: - 3.0 to 49.2%; pd: 95.6%). This effect was only present when controlling for maternal kinship, suggesting that groupmates or neighbours who are not maternally related may avoid synchronisation.

Since genetic kinship (i.e. combined maternal and paternal kinship) was known for only a subset of dyads, its inclusion as a predictor reduced the sample size by 52% (N = 432 litter combinations). Genetic kinship did not influence birth synchrony (estimate: 6.2%; 95% CI: −5.8 to 19.7%; pd = 82.2%), nor did presence within 50 meters in this subset model. Demographic maternal kinship and presence within two body lengths had similar effects on synchrony as reported in the previous model but were estimated with greater uncertainty (Supplementary Table S2B and Fig. S6).

### (b) The effect of kinship requires spatial proximity

To ensure that the effect of maternal kinship on synchrony did not merely reflect genetic similarity, we tested how this relationship changes with geographic distance between dens. This allowed us to determine whether maternal kinship synchronises females only when they live in the same area or if the effect persists across larger distances. We found that the synchronising effect of maternal kinship was restricted to mothers who lived nearby, whose dens were located within about 100 meters of each other (Fig. 1E, Supplementary Table S4 and Fig. S8). This suggests that the effect is socially mediated rather than driven by genetic similarity, as genetic effects would also apply at larger distances (but see Discussion).

### (c) Within-group birth synchrony increases with kinship but not relationship strength

Using fine-grained behavioural data from focal observations allowed us to explore the effect of differentiated social relationships beyond mere group membership and spatial association (Model 2). Within this smaller sample (N = 35 litter combinations), the time gap between births was 52.8% shorter for mother dyads with maternal kinship of 0.5 than for unrelated or unknown dyads (95% CI: -75.8 to -5.8%; pd: 98.2%), whereas no strong evidence for a synchronising effect was found for mother dyads with maternal kinship of 0.25 (estimate -37.5%; 95% CI: -70.5 to 36.3%; pd: 88.7%). Neither measure of relationship strength influenced synchrony (time spent within two body lengths: 95% CI: -36.9 to 46.2; pd: 57.1%; CSI: 95% CI: -44.0 to 29.7%; pd: 78.6%; Fig. 1F, Supplementary Table S5 an Fig. S9).

### (d) Allomaternal care increases with synchrony and maternal kinship

Using ad libitum observations, we examined the occurrence of allomaternal care among included litter dyads who belonged to the same group (descriptive data in Supplemental Text). The outcome variable was binary presence or absence of allomaternal care per litter combination (Model 3). We found that a smaller time gap between births (i.e., higher synchrony) increased the probability that at least one of the mothers allonursed the other litter (posterior mean log-odds estimate: -3.41; 95% CI: -5.91 to - 1.37, pd: 100%, Fig. 2A and C, Supplementary Table S6A). Independent of synchrony, allonursing was also more likely when maternal kinship between mothers was 0.5 (estimate: 2.80; 95% CI: -0.01 to 5.69; pd: 97.5%) compared with unrelated or unknown dyads. The same effect, but with higher uncertainty, was found for dyads with maternal kinship of 0.25 (estimate: 2.22; 95% CI: -0.95 to 5.35; pd: 91.9%). The probability of other forms of allomaternal care, i.e. allogrooming and lying in body contact, was similarly affected by synchrony and maternal kinship, but the effects were weaker (time gap estimate: -1.57; 95% CI: -3.94 to 0.20; pd: 95.8%; maternal kinship of 0.5 estimate: 2.87; 95% CI: -0.06 to 5.92; pd: 97.3%; maternal kinship of 0.25 estimate: 1.65; 95% CI: -1.57 to 4.97; pd: 84.8%, Fig. 2B and D).

**Fig 2.**
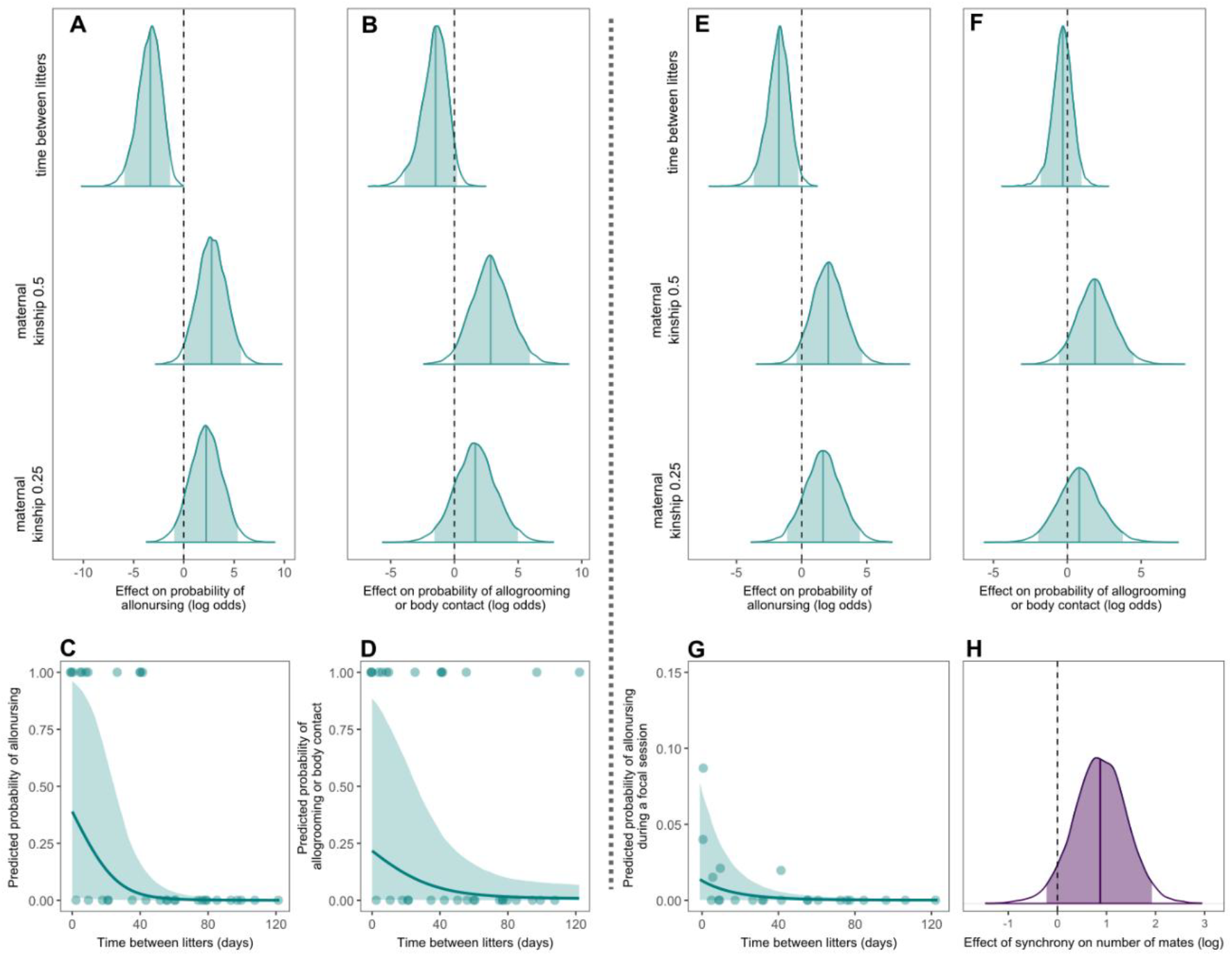
The effect of reproductive synchrony on allomaternal care and the number of mates.(**A-D**) Occurrence of allomaternal care per dyad, in function of synchrony and kinship. Results from Bayesian Bernoulli mixed-effects models examining the effects of birth synchrony (days between births) and maternal kinship (0.25 and 0.5, compared with non-kin) on the probability that allomaternal care occurs in a dyad (N = 33 litter combinations). Days between births were z-transformed (mean = 46.1, SD = 36.1). (**A, B**) Posterior distributions for the effects of synchrony and kinship on (**A**) allonursing and (**B**) on other forms of allomaternal care (allogrooming and lying in body contact). Solid lines: posterior means. Shaded areas: 95% credible intervals. Dashed line at zero: no effect. (**C, D**) Posterior predictions from the same models showing the effect of synchrony on (**C**) the probability of allonursing and (**D**) other forms of allomaternal care. Isolated effects are shown alongside raw data points, the line and shaded area show fitted values (posterior mean ± 95% CI). For the illustration, other predictors were held constant at their reference levels (maternal kinship = 0, data source = general) and random effects were excluded. (**E-G**) Frequency of allomaternal care in function of synchrony and kinship. Results from Bayesian binomial mixed-effects models examining the effects of birth synchrony (days between births) and maternal kinship (0.25 and 0.5, compared to non-kin) on the proportion of focal session with allomaternal care (N = 23 litter combinations). Days between births were z-transformed (mean = 45.0, sd = 36.9). (**E, F**) Posterior distributions for the effects of synchrony and kinship on (**E**) allonursing and (**F**) on other forms of allomaternal care (allogrooming and lying in body contact). Solid lines: posterior means. Shaded areas: 95% credible intervals. Dashed line at zero: no effect. (**G**) Posterior predictions from the same model as in (**E**) showing the effect of synchrony on the probability of allonursing. Isolated effects are shown alongside raw data points, the line and shaded area show fitted values (posterior mean ± 95% CI). For the illustration, maternal kinship was held constant at zero, random effects were excluded, and the y-axis was cropped at 0.15 due to low predicted probabilities. (**H**) Number of mates in function of oestrus synchrony. Posterior distribution of the Bayesian Poisson mixed-effects model, showing the effect of synchrony (number of fertile females in a group) on the number of males a focal female mated with on a given day. Solid lines: posterior means. Shaded areas: 95% credible intervals. Dashed line at zero: no effect.

Additional analysis of the frequency of allomaternal care during focal observations (Model 4) supports these results, showing that both synchrony and maternal kinship had a positive effect on allonursing frequency (Fig. 2E and G), whereas only close maternal kinship (0.5) was associated with higher frequencies of allogrooming and lying in body contact (Fig. 2F) (full results in Supplementary Text, Supplementary Table S6B).

### (e) Oestrus synchrony may reduce monopolisation by males

Finally, we used focal data on 8 females in heat to preliminarily explore the relationship between oestrus synchrony and monopolisation by males (Model 5). We found evidence that on days when multiple females from the same pack were simultaneously in heat, focal females were able to mate with more males (estimate on a log scale: 0.85, 95% CI: -0.25 to 1.90, pd: 94.2%, Fig. 2H). Indeed, females mated with multiple males only on days when another female was in heat, which then allowed them to mate with up to four males (Supplemental Fig. S9). In theory, this result could also be explained by a higher availability of males due to stronger attraction to two versus one oestrous female. However, in our study, two oestrous females did not attract more males per day (mean = 3.6, SD = 2.0) than a single female (mean = 3.7, SD = 1.9, Supplementary Fig. S9; visualisation of raw reproductive data for all included mothers and litters, see Supplementary Fig. S10).

## 4. Discussion

Our study provides evidence that female dogs synchronise giving birth, thereby increasing opportunities for allonursing in a population without strong reproductive seasonality. The results reveal that birth synchrony is higher among females that are more likely to cooperate: maternal relatives that live in the same area. This pattern suggests the presence of socially mediated synchronisation. We propose synchronisation to be adaptive by enabling cooperative breeding between females, and support this with our finding that more synchronous females and maternal relatives are indeed more likely to provide allomaternal care. Hence, synchronisation not only maintains systems where cooperation between breeding females is vital for offspring survival [39,40], but may also promote allomaternal care in facultative cooperative breeders. The preliminary finding that oestrous synchrony allows females to mate with more males suggests that synchronisation may confer multiple co-occurring benefits in polygynandrous species, perhaps contributing to the low rates of infanticide by male free-ranging dogs [54].

The higher birth synchrony we found among maternally related female dyads is consistent with reports showing that successful cooperative care in mammals most commonly occurs among kin [19,55–57]. According to our results, birth synchrony is only stronger among maternally related females when mothers breed at close distances. This indicates that synchronisation is socially mediated rather than purely determined by genetic timing mechanisms, as distance should not affect the latter. Supporting this argument, we found that whilst maternal kinship had a strong effect on synchrony, biparental kinship determined by genetic analysis did not. This indicates that synchronisation only occurs among related individuals who recognise each other as such, with paternal kin recognition limited or absent. Maternal care and early life interactions result in more reliable recognition of maternal versus paternal kin in mammals [67,68]. In dogs, this effect is likely reinforced by the polygynandrous mating system, which results in frequent occurrence of multiple paternity within litters and males fathering offspring in social groups other than their own [50].

Alternatively, the effect of genetic similarity on birth timing could be blurred if environmental differences result in phenotypic plasticity, i.e. one genotype produces different phenotypes under different environmental conditions. This could explain our results if distant females face larger environmental differences than immediate neighbours. However, our reported effect of close maternal kinship on synchrony vanished already at 100 m, where environmental differences are likely small.

Although maternal relatives need to belong to the same group to synchronise, additional increases in relationship strength beyond group membership did not translate into stronger synchronisation. The preferential synchronisation among maternally related females as opposed to unrelated females from the same group indicates that reproductive coordination occurs at the level of kin relationships rather than group membership alone. Hence, explanations based solely on shared environments and group membership are unlikely. Without knowing the underlying proximate mechanisms, we cannot explain why a stronger social relationship – and thus, higher exposure – does not translate into higher synchrony.

The link between synchronisation and cooperation is supported by our finding that, among mothers within the same group, both synchrony and maternal kinship independently increase the likelihood that allonursing and other forms of allomaternal care occur. Adding nuance by additionally examining differences in the frequency of caring behaviours, we found that synchrony and maternal kinship also increase the frequency of allonursing, whereas the frequency of other forms of allomaternal care is only higher among close maternal kin but not additionally influenced by synchrony. This shows that whilst maternally related females are more likely than unrelated females to provide any form of allomaternal care – confirming previous reports [49] – lactation’s physiological constraints render allonursing dependent on synchrony. These results support the key role of reproductive synchrony in facilitating allonursing and intensive communal care, as suggested by the strong birth synchrony found in vertebrates with obligate cooperative breeding among females, such as banded mongooses and greater anis [39,40], and in eusocial ants [69]. Strong synchrony among cooperatively breeding females can also be combined with litter mixing as a strategy to prevent infanticide, by hindering litter identification [39,40,54].

Unexpectedly, birth synchrony seemed reduced among mothers that live in the same area but are not close kin. This pattern emerged only after accounting for the effects of kinship, suggesting avoidance of synchrony among unrelated co-resident females. This may serve as a strategy to prevent competition and infanticide among females who would not cooperate during breeding. Without indirect benefits from allonursing related pups, strong asynchrony is more advantageous than synchrony as it prevents resource competition between litters. This argument is supported by a study on unrelated, communally nesting lab mice (*Mus musculus*), in which males can induce oestrus synchrony (Whitten and Bruce effects [27,70]). There, the risk of infanticide was lowest when females bred either asynchronously (avoiding communal nesting and competition) or highly synchronously (preventing litter discrimination within the communal nest), with the highest risk at intermediate synchrony where litters belong to the same communal nest but were discriminable [70]. Importantly, neither that study nor ours demonstrates strategic use of these differences, calling for further investigation. If confirmed, such fine-scaled synchronisation strategies within groups would be difficult to explain by environmental cues alone.

Our results indicate that oestrus synchrony allows females to mate with multiple partners, in accordance with the priority of access model [10] and with empirical studies in primates [38] and carnivores [71]. In species with limited variation in gestation length, oestrus synchrony largely translates into birth synchrony. Synchronisation may therefore arise either through direct selection on both traits or because one emerges as a by-product of selection on the other. Our finding that maternally related females synchronise more suggests that especially kin benefit from synchronising, pointing to primary selection on allonursing and allomaternal care through increased indirect fitness [19,55]. In contrast, direct benefits of monopolisation avoidance, including ensured fertilisation and reduced infanticide by males through paternity confusion [12–14], could be more effectively achieved by synchronising oestrus indiscriminately with all females from the same group or area, suggesting that this is not the primary target of selection in domestic dogs.

Although we included eight social groups, providing broad coverage across the study population, our study is limited by the restricted number of focal females and the comparatively brief focal observation period. These limitations may have left a potential association between birth synchrony and within-group social relationship strength undetected and may have reduced our ability to precisely estimate the association between oestrus synchrony and male monopolisation. Further study of these aspects and of potential contrasting strategies among individuals living in the same area will help to draw robust conclusions. Additionally, we could not analyse fitness effects of allonursing, since causes of high pup mortality were mostly unknown and may have been human-related, as reported for other free-ranging dog populations [53].

In summary, our results show that birth synchrony is higher among co-resident maternal kin in a large population of free-ranging dogs. This pattern could be explained by socially mediated synchronisation among females who are likely to breed cooperatively, which is additionally supported by our finding that both higher synchrony and maternal relatedness were independently associated with a higher likelihood of alloparental care and with higher rates of allonursing. Although the underlying mechanisms are unclear, previous studies have suggested olfactory cues may mediate reproductive synchronisation in mammals [25,26].

Our findings contribute to the debate surrounding socially mediated reproductive synchronisation [2– 4,30,32], highlighting the possibility that cases remain undetected due to overlooking cooperative care as the major selection force. Additionally, our results may explain why convincing reports of synchronisation are absent in species that do not require birth synchrony to enable cooperative care, such as humans. Due to the advantages of cooperative care, humans are able to combine slow maturation with shortened interbirth intervals [57]. Hence, aligned pregnancies within groups of familiar females may be advantageous by enabling allonursing (occurrence discussed by [72]). However, human oestrous cycles are short and occur frequently enough relative to the nursing period to not require socially mediated synchronisation. Instead, they may be optimally aligned by e.g. adjusting the timing of conception. We propose that future studies on socially mediated reproductive synchronisation would benefit from taking the cooperative care perspective more into account, both by comparing species with and without cooperative breeding and by considering whether socially mediated synchronisation would be required to enable cooperative care in a certain species.

## Acknowledgments

We thank our team in the field for their dedication, hard work and support. During data collection for this study, the Morocco team included: Lina Underberg, Karoline Stock, Lilo Schober, Debora Prince, Fabio Pensalfini, Valentina Napolitano, Flavia Mercuri, Chiara Meneghetti, Camilla Mancassola, Magdelena Juskaite, Maartje Hooymans, Clémence Helleu, Jérémy Hardouin, Léo Hanon, Claire Giraudet, Fiona Gaumard, Manon Delaunay, Giulia Cecchinato, Svenja Capitain, Maïwen Braconnier, Haytem Bouchri, Urša Blenkuš, Giulia Bigazzi, Anaïs Baillet and Corisande Abiven. Many thanks to Roger Mundry and Christof Neumann for their statistical advice. Shukran to HGVM for his support.

## Data and materials availability

Source data and R-code for statistics and figures in the paper will be made permanently accessible upon acceptance.

## Author Contributions

Conceptualisation: MV, NT, AB

Data curation: MV, NT, BC

Investigation: MV, NT, BC

Formal analysis: MV, NT, AB, CC, MP

Funding acquisition: AB, MP, SMP, FR, GC

Methodology: MV, AB, NT, GC, SMP, MP

Project administration: AB, GC, SMP, FR

Resources: FR

Supervision: AB Writing—original draft: MV

Writing—review & editing: MV, AB, FR, SMP, NT, BC, GC, CC, MP

## Ethics Statement

Ethical approval for DNA sampling was obtained from the Ethical Committee for Animal Veterinary Science and Public Health of the University Hassan II Rabat, Morocco (Protocol number: CESASPV_2023_04). All procedures applied in the present study were noninvasive and in accordance with the European Union Directive on the protection of animals used for scientific purposes (EU Directive 2010/63/EU). For behavioral observations, the Research Ethics Committee of the University of Veterinarian Medicine Vienna has confirmed that no ethical approval is required.

Observations were conducted under permission of the local authorities and in accordance with the ASAB guidelines for the treatment of animals in behavioral research (ASAB 2023).

## References

1. McClintock MK. 1971 Menstrual synchrony and suppression. Nature 229, 244–245. (doi:10.1038/229244a0)

2. Schank JC. 2001 Measurement and cycle variability: Reexamining the case for ovarian-cycle synchrony in primates. Behavioural Processes 56, 131–146. (doi:10.1016/s0376-6357(01)00194-2)

3. Schank JC. 2000 Can pseudo entrainment explain the synchrony of estrous cycles among golden hamsters (Mesocricetus auratus)? Hormones and Behavior 38, 94–101. (doi:10.1006/hbeh.2000.1603)

4. Wilson H. 1992 A critical review of menstrual synchrony research. Psychoneuroendocrinology 17, 565–591. (doi:10.1016/0306-4530(92)90016-z)

5. Fahs B, Gonzalez J, Coursey R, Robinson-Cestaro S. 2014 Cycling Together: Menstrual Synchrony as a Projection of Gendered Solidarity. Women’s Reproductive Health 1, 90–105. (doi:10.1080/23293691.2014.966029)

6. Lincoln GA, Clarke IJ, Hut RA, Hazlerigg DG. 2006 Characterizing a mammalian circannual pacemaker. Science 314, 1941–1944. (doi:10.1126/science.1132009)

7. van Schaik CP, Brockman DK. 2005 Seasonality in primate ecology, reproduction, and life history: An overview. In Seasonality in primates: Studies of living and extinct human and non-human primates (eds CP van Schaik, DK Brockman), pp. 3–20. Cambridge: Cambridge University Press.

8. Heideman PD, Utzurrum RC. 2003 Seasonality and synchrony of reproduction in three species of nectarivorous Philippines bats. BMC Ecology 3, 11. (doi:10.1186/1472-6785-3-11)

9. Pereira ME. 1991 Asynchrony within estrous synchrony among ringtailed lemurs (Primates: Lemuridae). Physiology & Behavior 49, 47–52. (doi:10.1016/0031-9384(91)90228-G)

10. Altmann S. 1962 A field study of the sociobiology of rhesus monkeys, Macaca mulatta. Annals of the New York Academy of Sciences 102, 338–435. (doi:10.1111/j.1749-6632.1962.tb13650.x)

11. Ostner J, Nunn CL, Schülke O. 2008 Female reproductive synchrony predicts skewed paternity across primates. Behav Ecol 19, 1150–1158. (doi:10.1093/beheco/arn093)

12. Hrdy SB. 2000 The optimal number of fathers: Evolution, demography, and history in the shaping of female mate preferences. Ann N Y Acad Sci 907, 75–96. (doi:10.1111/j.1749-6632.2000.tb06617.x.)

13. Hrdy SB. 1979 Infanticide among animals: A review, classification, and examination of the implications for the reproductive strategies of females. Ethology and Sociobiology 1, 13–40. (doi:10.1016/0162-3095(79)90004-9)

14. van Schaik CP, Hodges JK, Nunn CL. 2000 Paternity confusion and the ovarian cycles of female primates. In Infanticide by Males and its Implications (eds CP van Schaik, CH Janson), pp. 361–387. Cambridge University Press.

15. Ridley M. 1986 The number of males in a primate troop. Animal Behaviour 34, 1848–1858. (doi:10.1016/s0003-3472(86)80271-8)

16. Hasson O, Stone L. 2009 Male infertility, female fertility and extrapair copulations. Biological Reviews 84, 225–244. (doi:10.1111/j.1469-185X.2008.00068.x)

17. Santos RG, Pinheiro HT, Martins AS, Riul P, Bruno SC, Janzen FJ, Ioannou CC. 2016 The anti-predator role of within-nest emergence synchrony in sea turtle hatchlings. Proc. R. Soc. B. 283, 20160697. (doi:10.1098/rspb.2016.0697)

18. Boinski S. 1987 Birth synchrony in squirrel monkeys (Saimiri oerstedi). Behav Ecol Sociobiol 21, 393–400. (doi:10.1007/BF00299934)

19. Hayes LD. 2000 To nest communally or not to nest communally: A review of rodent communal nesting and nursing. Animal Behaviour 59, 677–688. (doi:10.1006/anbe.1999.1390)

20. De Oliveira Terceiro FE, Burkart JM. 2019 Cooperative Breeding. In Encyclopedia of Animal Cognition and Behavior (eds J Vonk, T Shackelford), pp. 1–6. Cham: Springer International Publishing. (doi:10.1007/978-3-319-47829-6_1351-1)

21. Ben Mocha Y, Scemama De Gialluly S, Griesser M, Markman S. 2023 What is cooperative breeding in mammals and birds? Removing definitional barriers for comparative research. Biological Reviews 98, 1845–1861. (doi:10.1111/brv.12986)

22. Chandra V, Fetter-Pruneda I, Oxley PR, Ritger AL, McKenzie SK, Libbrecht R, Kronauer DJC. 2018 Social regulation of insulin signaling and the evolution of eusociality in ants. Science 361, 398–402. (doi:10.1126/science.aar5723)

23. Burkart JM, van Schaik CP. 2010 Cognitive consequences of cooperative breeding in primates? Anim Cogn 13, 1–19. (doi:10.1007/s10071-009-0263-7)

24. Ims RA. 1990 The ecology and evolution of reproductive synchrony. Trends Ecol Evol 5, 135–140. (doi:10.1016/0169-5347(90)90218-3)

25. Sillero-Zubiri C, Johnson PJ, MacDonald DW. 1998 A hypothesis for breeding synchrony in Ethiopian wolves (Canis simensis). Journal of Mammalogy 79, 853. (doi:10.2307/1383093)

26. Berger J. 1992 Facilitation of reproductive synchrony by gestation adjustment in gregarious mammals: A new hypothesis. Ecology 73, 323–329. (doi:10.2307/1938743)

27. Bruce HM. 1959 An Exteroceptive Block to Pregnancy in the Mouse. Nature 184, 105–105. (doi:10.1038/184105a0)

28. Handelmann G, Ravizza R, Ray WJ. 1980 Social dominance determines estrous entrainment among female hamsters. Hormones and Behavior 14, 107–115. (doi:10.1016/0018-506X(80)90002-1)

29. McClintock MK. 1978 Estrous synchrony and its mediation by airborne chemical communication (Rattus norvegicus). Hormones and Behavior 10, 264–276. (doi:10.1016/0018-506X(78)90071-5)

30. Packer C, Pusey AE. 1983 Male takeovers and female reproductive parameters: A simulation of oestrous synchrony in lions (Panthera leo). Animal Behaviour 31, 334–340. (doi:10.1016/S0003-3472(83)80051-7)

31. French JA, Stribley JA. 1987 Synchronization of ovarian cycles within and between social groups in golden lion tamarins (Leontopithecus rosalia). American J Primatol 12, 469–478. (doi:10.1002/ajp.1350120403)

32. Clements MN, Clutton-Brock TH, Albon SD, Pemberton JM, Kruuk LEB. 2011 Gestation length variation in a wild ungulate. Functional Ecology 25, 691–703. (doi:10.1111/j.1365-2435.2010.01812.x)

33. Ihara Y, Collins DA, Oda R, Matsumoto-Oda A. 2016 Testing socially mediated estrous synchrony or asynchrony in wild baboons. Behav Ecol Sociobiol 70, 1921–1930. (doi:10.1007/s00265-016-2198-8)

34. Matsumoto-Oda A et al. 2007 Estrus cycle asynchrony in wild female chimpanzees, Pan troglodytes schweinfurthii. Behav Ecol Sociobiol 61, 661–668. (doi:10.1007/s00265-006-0287-9)

35. Strassmann BI. 1997 The biology of menstruation in Homo Sapiens: Total lifetime menses, fecundity, and nonsynchrony in a natural-fertility population. Current Anthropology 38, 123–129. (doi:10.1086/204592)

36. Tobler R, Pledger S, Linklater W. 2010 No evidence for ovarian synchrony or asynchrony in hamadryas baboons. Animal Behaviour 80, 829–837. (doi:10.1016/j.anbehav.2010.07.018)

37. Zerbe P, Clauss M, Codron D, Bingaman Lackey L, Rensch E, Streich JW, Hatt J, Müller DWH. 2012 Reproductive seasonality in captive wild ruminants: Implications for biogeographical adaptation, photoperiodic control, and life history. Biological Reviews 87, 965–990. (doi:10.1111/j.1469-185X.2012.00238.x)

38. Fürtbauer I, Mundry R, Heistermann M, Schülke O, Ostner J. 2011 You mate, I mate: Macaque females synchronize sex not cycles. PLOS ONE 6, e26144. (doi:10.1371/journal.pone.0026144)

39. Hodge SJ, Bell MBV, Cant MA. 2011 Reproductive competition and the evolution of extreme birth synchrony in a cooperative mammal. Biol. Lett. 7, 54–56. (doi:10.1098/rsbl.2010.0555)

40. Riehl C. 2016 Infanticide and within-clutch competition select for reproductive synchrony in a cooperative bird. Evolution 70, 1760–1769. (doi:10.1111/evo.12993)

41. Gilchrist JS. 2006 Female eviction, abortion, and infanticide in banded mongooses (Mungos mungo): implications for social control of reproduction and synchronized parturition. Behavioral Ecology 17, 664–669. (doi:10.1093/beheco/ark012)

42. Lord K, Feinstein M, Smith B, Coppinger R. 2013 Variation in reproductive traits of members of the genus Canis with special attention to the domestic dog (Canis familiaris). Behavioural Processes 92, 131–142. (doi:10.1016/j.beproc.2012.10.009)

43. Macdonald DW, Campbell LAD, Kamler JF, Marino J, Werhahn G, Sillero-Zubiri C. 2019 Monogamy: cause, consequence, or corollary of success in wild canids? Frontiers in Ecology and Evolution 7. (doi:10.3389/fevo.2019.00341)

44. Moehlman PD, Hofer H. 1996 Cooperative breeding, reproductive suppression, and body mass in canids. In Cooperative breeding in mammals (eds NG Solomon, JA French), pp. 76–128. Cambridge University Press. (doi:10.1017/CBO9780511574634.005)

45. Asa CS. 1996 Hormonal and experiential factors in the expression of social and parental behavior in canids. In Cooperative breeding in mammals (eds NG Solomon, JA French), pp. 129–149. Cambridge University Press. (doi:10.1017/CBO9780511574634.006)

46. Randall DA, Pollinger JP, Wayne RK, Tallents LA, Johnson PJ, Macdonald DW. 2007 Inbreeding is reduced by female-biased dispersal and mating behavior in Ethiopian wolves. Behavioral Ecology 18, 579–589. (doi:10.1093/beheco/arm010)

47. Cafazzo S, Valsecchi P, Bonanni R, Natoli E. 2010 Dominance in relation to age, sex, and competitive contexts in a group of free-ranging domestic dogs. Behavioral Ecology 21, 443–455. (doi:10.1093/beheco/arq001)

48. Daniels TJ. 1983 The social organization of free-ranging urban dogs. II. Estrous groups and the mating system. Applied Animal Ethology 10, 365–373. (doi:10.1016/0304-3762(83)90185-2)

49. Paul M, Bhadra A. 2018 The great Indian joint families of free-ranging dogs. PLoS ONE 13, e0197328. (doi:10.1371/journal.pone.0197328)

50. Car C et al. 2025 Mating system of free-ranging domestic dogs and its consequences for dog evolution. Proc. Natl. Acad. Sci. U.S.A. 122, e2421756122. (doi:10.1073/pnas.2421756122)

51. Fielding HR, Gibson AD, Gamble L, Fernandes KA, Airikkala-Otter I, Handel IG, Bronsvoort BM de C, Mellanby RJ, Mazeri S. 2021 Timing of reproduction and association with environmental factors in female free-roaming dogs in southern India. Preventive Veterinary Medicine 187, 105249. (doi:10.1016/j.prevetmed.2020.105249)

52. Pal SK, Roy S, Ghosh B. 2021 Pup rearing: The role of mothers and allomothers in free-ranging domestic dogs. Applied Animal Behaviour Science 234, 105181. (doi:10.1016/j.applanim.2020.105181)

53. Pal SK. 2005 Parental care in free-ranging dogs, Canis familiaris. Applied Animal Behaviour Science 90, 31–47. (doi:10.1016/j.applanim.2004.08.002)

54. Vanderheyden M, Chaignon B, Car C, Pilot M, El Berbri I, Marshall‐Pescini S, Range F, Berghänel A. 2025 A direct observation of infanticide by a female free‐ranging dog (Canis familiaris) supports the resource competition hypothesis. Ecology and Evolution 15, e72727. (doi:10.1002/ece3.72727)

55. Packer C, Lewis S, Pusey A. 1992 A comparative analysis of non-offspring nursing. Animal Behaviour 43, 265–281. (doi:10.1016/S0003-3472(05)80222-2)

56. Roulin A. 2002 Why do lactating females nurse alien offspring? A review of hypotheses and empirical evidence. Animal Behaviour 63, 201–208. (doi:10.1006/anbe.2001.1895)

57. Hrdy SB. 2005 Evolutionary context of human development: The cooperative breeding model. In Attachment and bonding: a new synthesis (eds SC Carter, L Ahnert, K Grossmann, SB Hrdy, ME Lamb, SW Porges, N Sachser), pp. 9–32. Cambridge, MA: MIT Press.

58. Riehl C, Strong MJ. 2018 Stable social relationships between unrelated females increase individual fitness in a cooperative bird. Proc. R. Soc. B. 285, 20180130. (doi:10.1098/rspb.2018.0130)

59. Berghänel A, Lazzaroni M, Ferenc M, Pilot M, El Berbri I, Marshall-Pescini S, Range F. 2025 Cofeeding at rich clumped food patches in free-ranging dogs: social tolerance or scramble competition? Behav Ecol Sociobiol 79, 51. (doi:10.1007/s00265-025-03590-8)

60. Range F, Marshall-Pescini S. 2022 Wolves and dogs: between myth and science. Cham: Springer International Publishing.

61. Pilot M et al. 2015 On the origin of mongrels: evolutionary history of free-breeding dogs in Eurasia. Proc. R. Soc. B. 282, 20152189. (doi:10.1098/rspb.2015.2189)

62. Altmann J. 1974 Observational study of behavior: Sampling methods. Behav 49, 227–266. (doi:10.1163/156853974×00534)

63. Schülke O et al. 2022 Quantifying within-group variation in sociality—covariation among metrics and patterns across primate groups and species. Behav Ecol Sociobiol 76, 50. (doi:10.1007/s00265-022-03133-5)

64. Bürkner P-C. 2017 brms: An R package for Bayesian multilevel models using Stan.

65. R Core Team. 2025 R: A language and environment for statistical computing. Vienna, Austria: R Foundation for Statistical Computing. See https://www.R-project.org/.

66. Fox J, Weisberg S. 2019 An R companion to applied regression.

67. Smith JE. 2014 Hamilton’s legacy: Kinship, cooperation and social tolerance in mammalian groups. Animal Behaviour 92, 291–304. (doi:10.1016/j.anbehav.2014.02.029)

68. Hepper PG. 1994 Long-term retention of kinship recognition established during infancy in the domestic dog. Behavioural Processes 33, 3–14. (doi:10.1016/0376-6357(94)90056-6)

69. Piqueret B, Weissflog J, Tretter S, Zetzsche T, Veit D, Bartram S, Halitschke R, Ulrich Y. 2026 Offspring chemical control of adult reproductive transitions in a social insect. Proc. Natl. Acad. Sci. U.S.A. 123, e2526776123. (doi:10.1073/pnas.2526776123)

70. Schmidt J, Kosztolányi A, Tökölyi J, Hugyecz B, Illés I, Király R, Barta Z. 2015 Reproductive asynchrony and infanticide in house mice breeding communally. Animal Behaviour 101, 201–211. (doi:10.1016/j.anbehav.2014.12.015)

71. Say L, Pontier D, Natoli E. 2001 Influence of oestrus synchronization on male reproductive success in the domestic cat (Felis catus L.). Proceedings of the Royal Society of London. Series B: Biological Sciences 268, 1049–1053. (doi:10.1098/rspb.2001.1583)

72. Hewlett BS, Winn S. 2014 Allomaternal nursing in humans. Current Anthropology 55, 200–229. (doi:10.1086/675657)

