## Supplementary Material for "Birth synchronisation increases allonursing opportunities in free-ranging dogs"

This PDF file includes:

Supporting text
Figures S1 to S12
Tables S1 to S6
SI References

Other supporting materials for this manuscript include the following:

Datasets S1 to S5
Scripts S1 to S5

Supporting Information Text

1. Extended Methods

a) Litter dyad composition

Litter dyads were composed of the temporally nearest parturition between two mothers (Supplementary Fig. S2). To ensure this, we only included dyads for which we knew a) the birth date of the previous litter of the mother with the later parturition date (unless she was primiparous) and b) the birth date of the subsequent litter of the mother with the earlier parturition date.

Missed pregnancies or unrecorded parturitions (e.g., due to immediate litter loss) could result in incorrectly linked litter dyads. In such cases, the recorded previous or subsequent birth may not have been the true adjacent event but rather the next-but-one, potentially leading to erroneous synchrony values. Hence, we only included litter combinations where the respective previous or following parturition occurred within 320 days. Since the shortest interbirth interval observed in our study population was 162 days (Supplementary Fig. S12; see also [1]), two consecutive births of the same female occurring within less than 320 days can be excluded, effectively ruling out misclassification of next-but-one births as adjacent events. To avoid extraordinarily large time gaps between parturitions, resulting from extended interbirth intervals that might reflect impaired reproductive performance and thus bias our results, we further applied an upper cutoff time of 140 days on the synchrony values. This threshold represents the time window within which two reproductively healthy females are likely to give birth (Supplementary Fig. S12 and [1]).

b) Maternal kinship from demographic data

Presence of maternal kinship was determined for 63 dyads by observing parturitions or dogs younger than 6 months being nursed or groomed by a single (lactating) female. Observations indicating maternal kinship were done ad libitum, starting in April 2022, and predominantly concerned newborn pups. We included maternal kinship as a factor with three categories of expected relatedness: 0.5 (mother-daughter), 0.25 (e.g. grandmother-granddaughter) and 0 (non-kin or unknown). Siblings were treated as half-siblings (0.25) since full-siblings could not be identified based on demographic data, given the occurrence of multiple paternity within litters [2]. Maternal kinship of less than 0.25 (e.g. cousins) was known for very few dyads but might have remained unrecognized for many others based on the available demographic data. Additionally, observations in primates show less altruism in dyads with low levels of relatedness compared to e.g. mother-offspring pairs [3,4]. This is likely due, at least in part, to the animals' difficulty of identifying more distant kin [5,6]. Hence, we classified all dyads with kinship <0.25 as 'non-kin'. Notably, this more conservative approach could raise type two but not type one errors in our analyses. Among dyads classified as mother–daughters based on behavioural observations, genetic data were available for 44% of cases and corroborated the classification in all but one instance.

c) Kinship from genetic data

Saliva samples were collected from 57 out of the 64 individuals included in this report, using non-invasive collection kits (Performagene PG 100, DNA Genotek, Canada). The DNA was extracted according to the manufacturer's instructions. SNP genotyping was conducted using the Axiom Canine HD Genotyping Array (Thermo Scientific). For 37 individuals, DNA extraction and genotyping were successful, generating a dataset of 201,723 autosomal SNPs with less than 10% missing data. Plink1.9 software was used for data filtering [7] and to estimate relatedness using the pair-wise identity by descent (IBD) coefficient pi-hat. This was done for 342 included mother dyads. For a full description of the genetic kinship reconstruction, see [8].

d) Social relationship strength calculation

Proximity data yielded high proportions of zeroes (Fig. 1C) since dyads were linked across the population and individuals lived up to 5 km apart. Because including these data as a continuous predictor led to zero-inflation, we instead included relationship strength as a binary measure (whether two individuals were seen within a certain distance but not considering how often). Since reproductive synchronisation likely occurs during the months before conception, we calculated these proximities for the three months preceding the earlier of the two estimated conception dates in each dyadic litter combination. Conception dates were calculated by subtracting 62 days, the average gestation length in dogs [1], from the parturition date.

Using the focal data, dyadic relationship strength was calculated based on 1) proximity and 2) composite sociality index (CSI). The proximity measure was the time spent within two body lengths, divided by the summed observation time of both individuals. The CSI [9] was composed using the frequency and/or duration of the following affiliative behaviours: body contact, grooming, play, greeting and friendly postures, corrected by the summed focal observation time of both individuals.

$$\mathrm{CSI}_{xy} = \frac{\sum_{i=1}^{d} \frac{f_{ixy}}{\overline{f_i}}}{d}$$

with d the number of behaviours that contribute to the index, $f_{ixy}$ the rate of behaviour i for dyad xy, and $\overline{f_i}$ the mean rate of behaviour i across all dyads in the subset of interest (in this study: all included females). The rate $f_{ixy}$ of each behaviour is calculated by dividing the total duration or count of the behaviour by the summed focal observation time of both partners x and y. The behaviours included in the CSI were: grooming (duration and count), play (duration and count), body contact (duration and count), greetings (count) and friendly postures (count).

Since focal data were collected with higher density than the scan data, we calculated relationship strength based on the data collected for one month (instead of three) preceding the earliest estimated conception date within the dyad, allowing us to include more dyads. In dyads for which both values were available, they were strongly correlated (Spearman's rank correlation for proximity: rho = 0.84, $p < 0.001$; CSI: rho = 0.78, $p < 0.001$). In both the scan and the focal data set, dyads were only included if both mothers were present in the study area during the respective observation period.

Group membership was determined based on social network measures derived from the scan observations, using the Walktrap algorithm in igraph [10] (see below). Only focal dyads from the same social group were included. Affiliative behaviour never occurred between focal females from different groups.

#### e) Walktrap algorithm for group determination

Social networks were constructed using data from scan observations, carried out between April 16th, 2022, and December 31st, 2024. Edges were constructed based on the number of times dogs were seen within five meters of each other, after correcting these counts for the number of scans in which at least one of the two individuals was observed during the time when their general presence in the study area overlapped (Simple Ratio Index, [11]). The correction accounted for demographic changes, missed dyadic interactions outside the scan area and differences in sampling frequency. An individual's general presence in the study area was defined by its first and last observation. Community structure was detected using the Walktrap algorithm in igraph [10], applied after including only the strongest 75% of connections to reflect stable relationships. This cutoff was necessary given the significant flexibility in social organization displayed by free-ranging dogs. Whilst many dogs are part of social groups, mixing with non-group members was common, especially within the large time window we used to build the social network.

#### f) Allonursing frequency

Model 4 investigated the effects of kinship and synchrony on the frequency of allomaternal care and allonursing within focal groups recorded during the focal observations. Whilst frequency measures add more nuance compared to the analysis of occurrence per dyad, these were only available for focal females, limiting the sample size (N = 23 litter combinations with 22 litters from 17 females). These included all instances of focal individuals providing allomaternal care and of allomaternal care being provided to a focal female's pups within a given litter dyad. Although the latter type of observation is ad libitum rather than focal (since neither the allomother nor the litter are the focal individual), mothers generally spent most of their time within view of their litter during the first three months after parturition, allowing us to record all allomaternal care which occurred during a focal observation of the mother with high accuracy. Allomaternal care was recorded as a binary outcome per focal session and modelled as a proportion of focal sessions per dyad, with between 43 and 176 included focal sessions per litter dyad. A binomial model with a logit link function was used, specifying the number of underlying focal sessions per litter combination as the number of trials. The same priors as in Model 3 were used.

### 2. Extended Results

#### a) Allonursing occurrence: descriptive data

Of the 42 litters included in this analysis, 13 (31%) were allonursed and 17 (40%) received other forms of allomaternal care (grooming or lying in body contact). Litters received allomaternal care from older (half-) sisters (41% of allonursed litters; 38% of litters receiving grooming or body contact), grandmothers (35% and 33%, respectively), (half-) aunts (12% and 14%), and unrelated females (12% and 14%). In one instance, a mother-daughter pair gave birth in the same den within hours of each other. When only considering the 21 litters born to focal females and observed more intensely, we found higher percentages of litters that were seen to receive allonursing (12 litters; 57%) or other forms of allomaternal care (17 litters; 81%).

#### b) Model 4: Allonursing frequency

Analysing the effects of synchrony and maternal kinship on the frequency of allomaternal care (Model 4, Bayesian binomial mixed-effects model, N = 23 litter combinations from 17 mothers), we found that the frequency of allonursing increased with decreasing time between births (estimate: -1.79; 95% CI: -3.63 to -0.23; pd: 99.0%) and with increasing maternal kinship. The evidence that maternal kinship of 0.5 increased the frequency of allonursing among mother dyads was strong (estimate: 2.09; 95% CI: -0.35 to 4.64; pd: 95.4%), but weaker for maternal kinship of 0.25 (estimate: 1.67; 95% CI: -1.12 to 4.49; pd: 88.5%) compared to unrelated or unknown dyads (Fig. 2E and G, Supplementary Table S6B). Only maternal kinship of 0.5 showed a positive effect on the frequency of allogrooming and lying in body contact (estimate: 1.89; 95% CI: -0.51 to 4.47; pd: 94.0%) (Fig. 2F), whereas maternal kinship of 0.25 (estimate: 0.84; 95% CI: -1.98 to 3.76; pd: 72.1%) and birth synchrony (estimate: -0.33; 95% CI: -1.79 to 0.95, pd: 68.9%) showed no clear association.

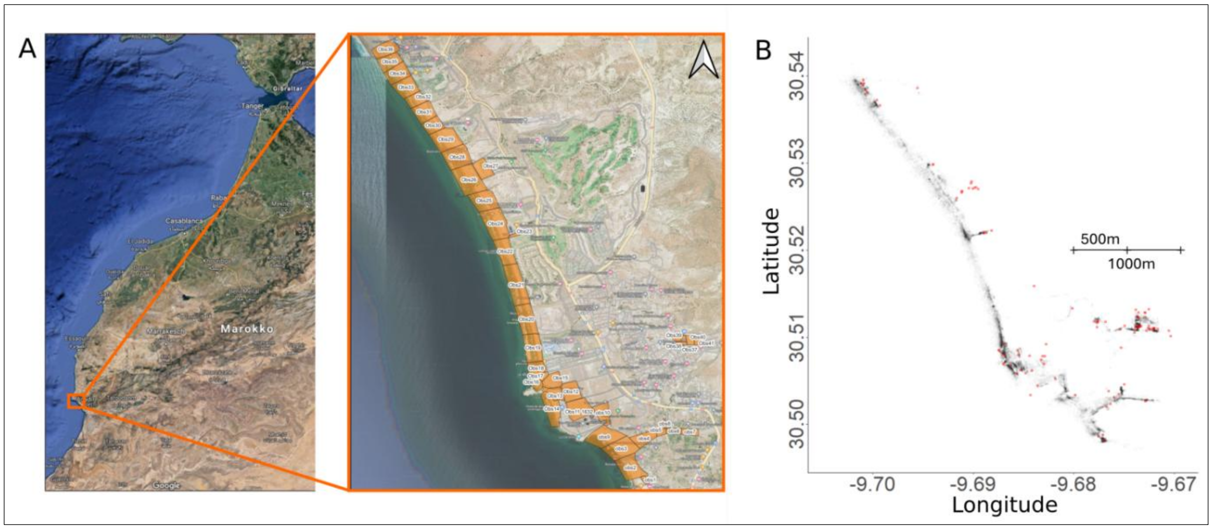

Fig. S1

Study area and the distribution of dogs and dens. (**A**) Study site with the areas of regular data collection. (**B**) All GPS positions of individual dogs recorded during the scan observations between 2022 and 2024 (grey) and all GPS positions of dens recorded in the same period (red).

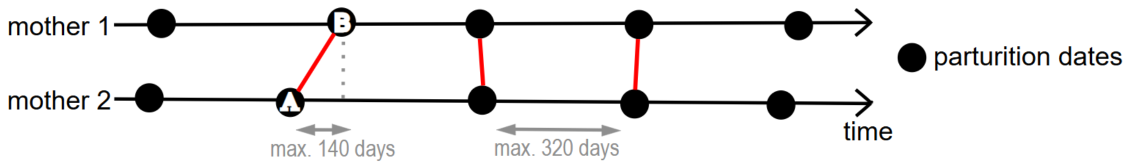

Fig. S2

Illustration of how parturition dates of two mothers were linked dyadically. Each litter from mother 1 was paired with only a single litter from mother 2, specifically the litter closest in time. The number of days between the linked births was used as a measure of reproductive synchrony, with larger values indicating lower synchrony.

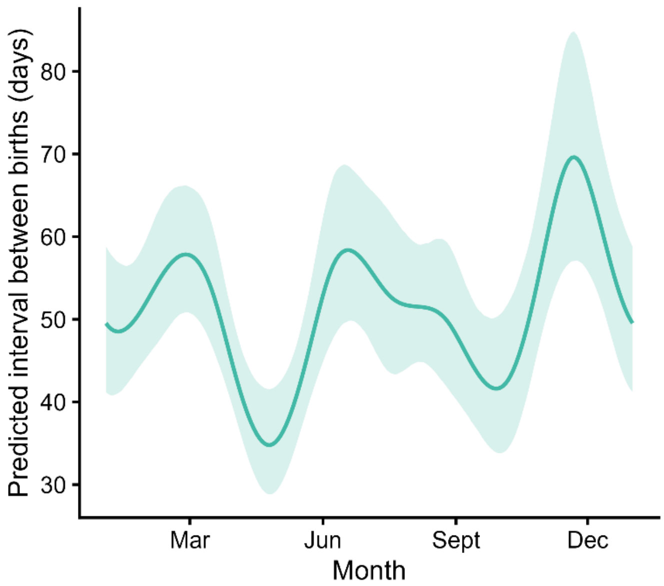

Fig. S3

Visualisation of predicted seasonal effect of day on the time between births (posterior means ± 95% CI) from the Bayesian Gamma mixed-effects model (Model 1). Seasonality was modelled as a cyclic smooth function of day, with day 1 referring to January 1st. For the illustration, other predictors were held constant at their reference level (maternal kinship = 0, relationship strength = 0) and random effects were excluded.

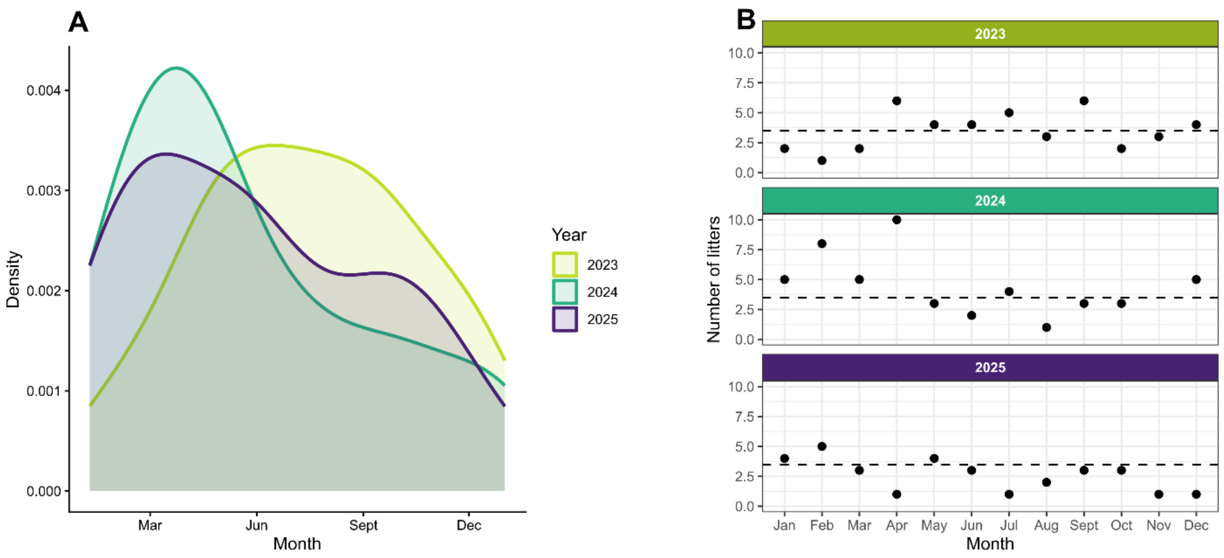

Fig. S4

Distribution of litter births throughout the year, visualised as (A) a density plot with ‘day 1’ referring to 1 January and (B) the raw number of litters per month, shown separately for each year. The dotted line shows the average number of litters per month, across the three years. All litters recorded in 2023, 2024 and 2025 for which parturition dates were estimated with an estimation error of max. ± 7 days were included (N = 122 litters). Only years with complete year-round data collection were selected.

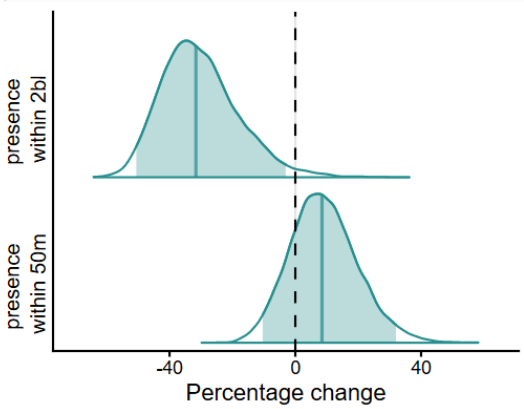

Fig. S5

Reduced Model 1 - Posterior distributions from the reduced Bayesian Gamma mixed-effects model, showing the isolated effect of relationship strength (based scan proximities: presence within two body lengths and within 50 m) on the time between births. Except for the removal of kinship-related terms, the model was identical to the full model including all dyads (N = 898, Table S2A).

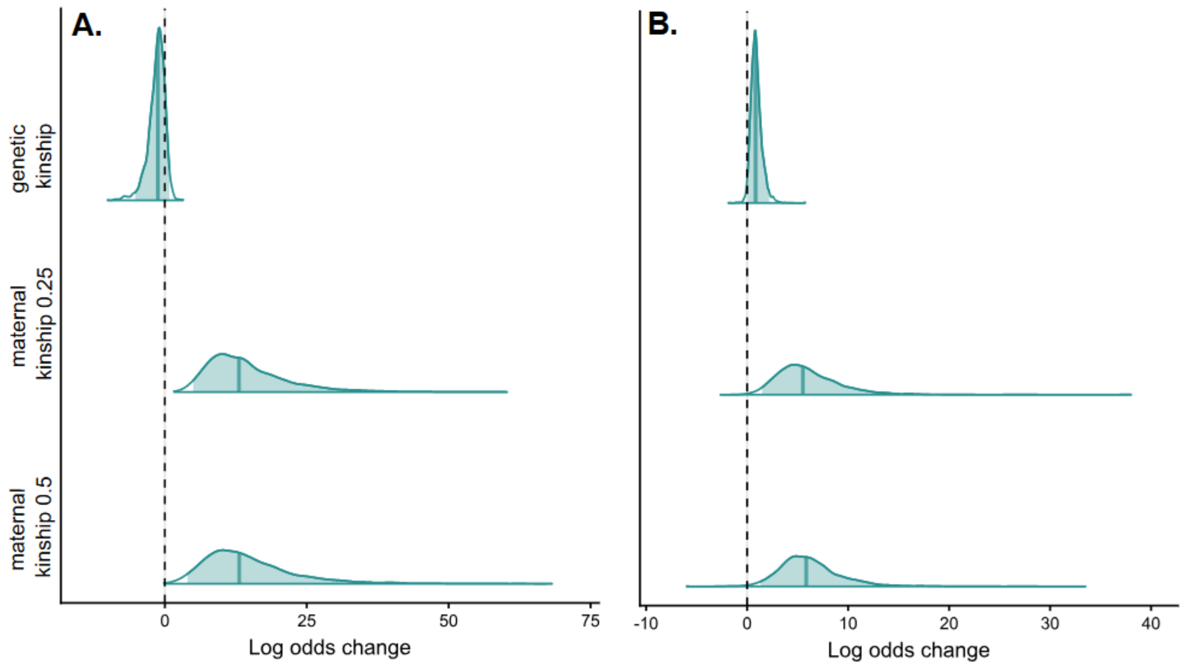

Fig. S6

Posterior distributions from the Bayesian Bernoulli mixed-effects models testing the effect of maternal and genetic kinship on the probability of (A) being seen within two body lengths and (B) within fifty meters. Estimates are given in log odds. Genetic kinship was z-transformed (mean = 0.06, sd = 0.1).

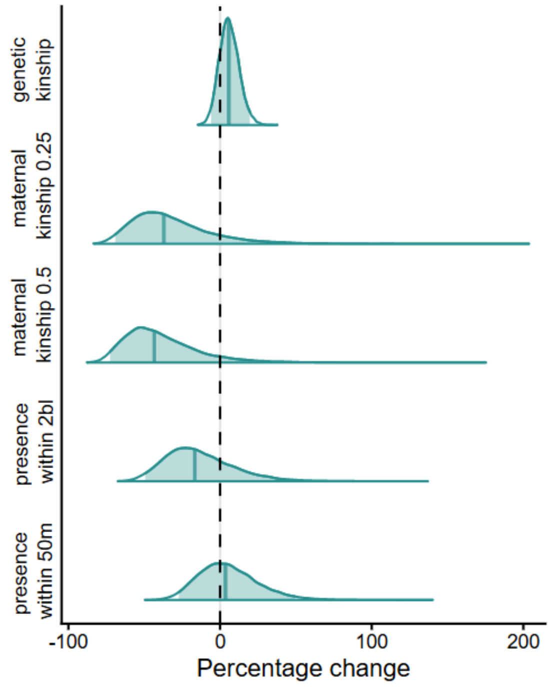

Fig. S7

Posterior distributions from Bayesian Gamma mixed-effects models showing the effect of relationship strength and kinship on the time between births, using scan proximities (within two body lengths and within 50 m, corrected according to the Simple Ratio Index, see Methods) to approximate relationship strength. Both maternal and genetic kinship were included as predictors (N = 432 litter combinations, Table S3B). Reference category for maternal kinship: non-kin or unknown. Genetic kinship coefficients were z-transformed (mean = 0.06, SD = 0.1). Full lines are posterior means, the shaded areas show the 95% credible intervals, and the dashed vertical line at zero indicates no effect.

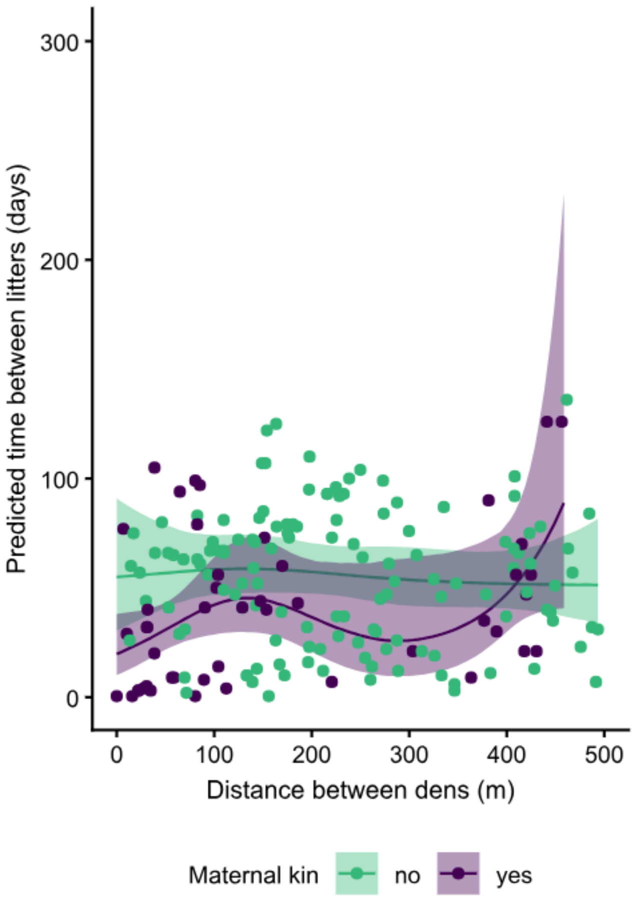

Fig. S8

Posterior predictions from the Bayesian Gamma mixed-effects model with separate smooth functions for kin and non-kin dyads, showing the time difference between litter births as a function of the distance between dens. Isolated predicted effects for kin versus non-kin, alongside raw data points. The solid lines show posterior means, shaded areas indicate 95% credible intervals. For the illustration, maternal kinship was held constant at zero and random effects were excluded.

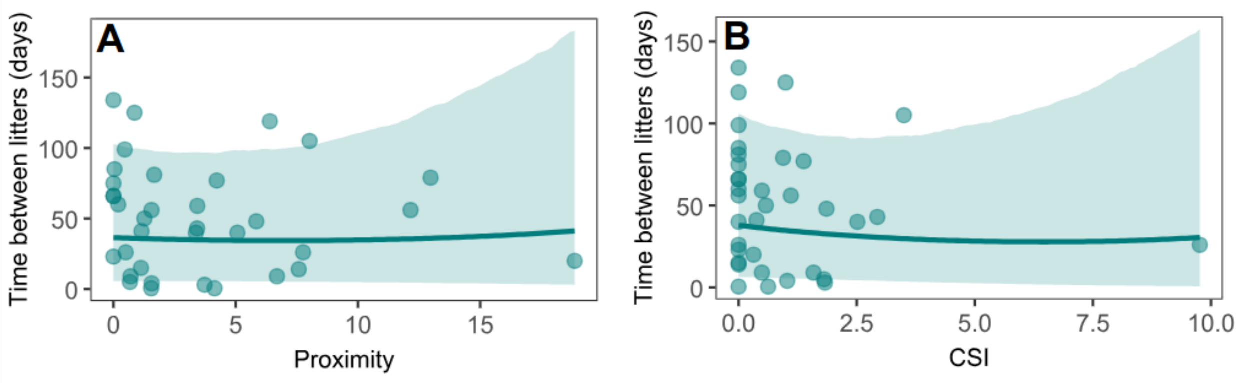

Fig. S9

Synchrony within social groups, in function of social relationship strength. Isolated effects of (A) proximity and (B) CSI on synchrony are shown alongside raw data points. The line and shaded area showing fitted values (posterior mean ± 95% CI). For the illustration, other predictors were held constant or at their reference levels (mean day = 169.2, maternal kinship = 0, mean CSI = 1.0, mean proximity = 3.6) and random effects were excluded.

Fig. S10.

Visualisation of raw reproductive data per individual. Rectangles represent days on which the individual was observed, colours indicate its reproductive state and grey indicating no visible signs. Full circles represent litters. Neutering events are shown for multiple days after the procedure to enhance visibility. All reproductive data of females included in this study are shown on the graph, including litters which did not meet the inclusion criteria (see Methods). Birth dates range from 22 September 2022 to 12 May 2025. Large boxes delineate social groups. Note that observation effort varied over time, with most intense data on group-living females collected during the year-long focal observation period, from March 2023 to March 2024.

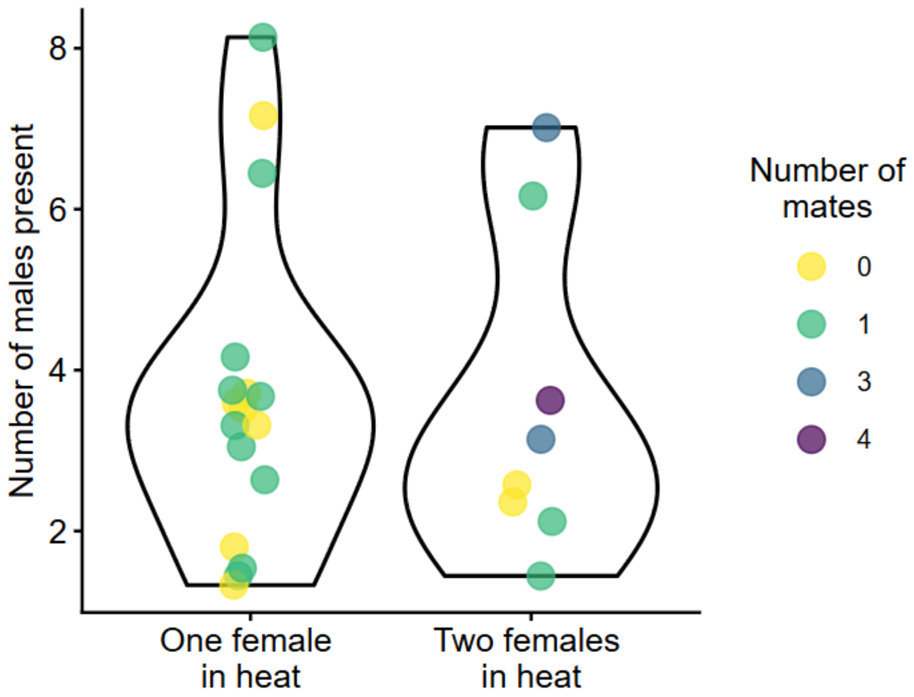

Fig. S11.

Daily mean number of males observed within 25 m of a focal female in heat when she was the only female in heat in the group versus when another female was simultaneously in oestrus. Colour indicates the number of males the focal female mated with on each day.

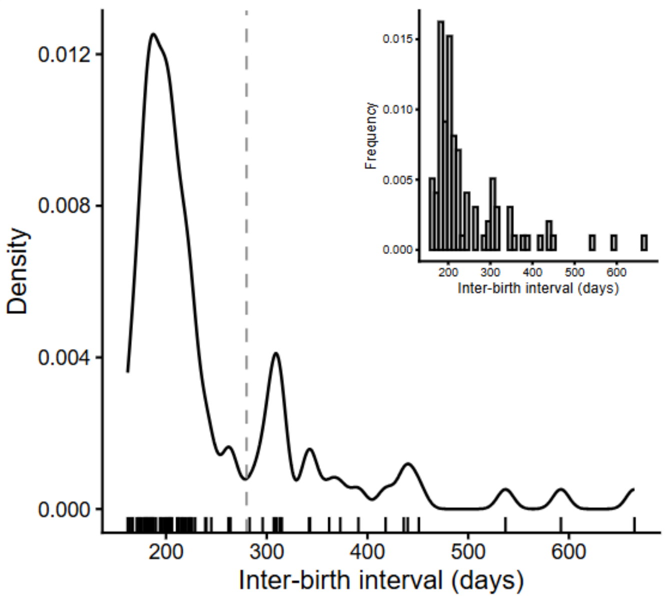

Fig. S12

Density function of inter-birth intervals (IBIs), showing which interval lengths are more likely. Raw data are shown as a rug plot along the x-axis (N = 96 IBIs). Inset: the respective histogram. The shortest IBI was 162 days (including IBIs following early litter loss), and the main peak including the majority of IBIs (75,0%) was below 280 days (dashed line). Assuming this value as a maximum IBI, two females with regular cycles can be expected to have their closest parturitions within 140 days of each other.

Table S1.

Reduced Model 1 - Output of the reduced Bayesian Gamma mixed-effects model, showing the isolated effect of relationship strength (presence within two body lengths and within 50 m) on the time between births. Except for the removal of kinship-related terms, the model was identical to Model 1 including all dyads (N = 898 litter combinations with 108 litters from 64 females, Table S2A).

| | Estimate | | 95% CI | | Probability of direction (%) |
|---|---|---|---|---|---|
| | Log scale | % Change | Log scale | % Change | |
| Intercept | 3.91 | - | 3.85 to 3.97 | - | - |
| Seen within 50 m | 0.08 | 8.3 | -0.1 to 0.28 | -9.5 to 32.3 | 79.9 |
| Seen within two body lengths | -0.38 | -31.6 | -0.71 to -0.03 | -50.8 to -3.0 | 98.4 |
| | SD | | 95% CI | | |
| Smooth term | | | | | |
| Seasonality | 0.24 | | 0.10 to 0.48 | | |
| Random effects | | | | | |
| Mother dyad | 0.04 | | 0.00 to 0.10 | | |
| Mother | 0.06 | | 0.00 to 0.18 | | |
| Litter | 0.08 | | 0.00 to 0.21 | | |

Table S2.

Model 1 - Output of the Bayesian Gamma mixed-effects models, showing the effect of relationship strength and kinship on the time between births, using scan proximities (two body lengths and 50 m) to approximate relationship strength. Genetic kinship was not included in (A) in order to maintain a larger sample size (N = 898 litter combinations with 108 litters from 64 females) but was included in (B), reducing the sample (N = 432 litter combinations with 75 litters from 39 females). The effects of maternal kinship categories 0.25 and 0.5 are in comparison with non-kin or dyads with unknown kinship. Genetic kinship coefficients were z-transformed (mean = 0.06, SD = 0.1).

| | A) Without genetic kinship (N = 898) | | | | |
|---|---|---|---|---|---|
| | Estimate | | 95% CI | | |
| | Log scale | % Change | Log scale | % Change | Probability of direction (%) |
| Intercept | 3.92 | - | 3.85 to 3.98 | - | - |
| Seen within 50 m | 0.19 | 20.9 | -0.03 to 0.40 | -3.0 to 49.2 | 95.6 |
| Seen within 2 body lengths | -0.27 | -23.7 | -0.61 to 0.09 | -45.7 to 9.4 | 93.3 |
| Maternal kinship 0.5 | -0.41 | -33.6 | -0.76 to -0.03 | -53.2 to -3.0 | 98.4 |
| Maternal kinship 0.25 | -0.25 | -22.1 | -0.57 to 0.08 | -43.4 to 8.3 | 93.1 |
| | SD | | 95% CI | | |
| Smooth term | | | | | |
| Seasonal | 0.23 | | 0.10 to 0.48 | | |
| Random effects | | | | | |
| Mother dyad | 0.04 | | 0.00 to 0.10 | | |
| Mother | 0.07 | | 0.00 to 0.18 | | |
| Litter | 0.09 | | 0.00 to 0.22 | | |
| | B) With genetic kinship (N = 432) | | | | |
| | Estimate | | 95% CI | | |
| | Log scale | % Change | Log scale | % Change | Probability of direction (%) |
| Intercept | 3.97 | - | 3.88 to 4.06 | - | - |
| Seen within 50 m | 0.04 | 4.1 | -0.32 to 0.4 | -27.4 to 49.2 | 57.5 |
| Seen within 2 body lengths | -0.18 | -16.5 | -0.68 to 0.33 | -49.3 to 39.1 | 75.9 |
| Maternal kinship 0.5 | -0.57 | -43.4 | -1.29 to 0.19 | -72.5 to 20.9 | 92.8 |
| Maternal kinship 0.25 | -0.46 | -36.9 | -1.17 to 0.29 | -69.0 to 33.6 | 88.8 |
| Genetic kinship | 0.06 | 6.2 | -0.06 to 0.18 | -5.8 to 19.7 | 82.2 |
| | SD | | 95% CI | | |
| Smooth term | | | | | |
| Seasonal | 0.17 | | 0.01 to 0.42 | | |
| Random effects | | | | | |
| Mother dyad | 0.06 | | 0.00 to 0.16 | | |
| Mother | 0.07 | | 0.00 to 0.21 | | |
| Litter | 0.08 | | 0.00 to 0.23 | | |

Table S3.

Output of the Bayesian Bernoulli mixed-effects models testing the effect of maternal and genetic kinship on the probability of being seen (A) within two body lengths (B) and within 50 m (N = 432 litter combinations with 75 litters from 39 females). The effects of maternal kinship categories 0.25 and 0.5 are in comparison with non-kin or dyads with unknown kinship. Random effects were identical to the other models. No priors were used. Estimates are given in log odds. Genetic kinship was z-transformed (mean = 0.06, sd = 0.1).

| | Estimate | 95% CI | Probability of direction (%) |
|---|---|---|---|
| A) Two body lengths | | | |
| Intercept | -9.49 | -20.43 to -5.05 | - |
| Maternal kinship 0.25 | 14.61 | 4.97 to 34.58 | 100 |
| Maternal kinship 0.5 | 14.81 | 3.97 to 36.27 | 99.98 |
| Genetic kinship | -1.47 | -5.24 to 0.78 | 87.10 |
| | SD | 95% CI | |
| Random effects | | | |
| Mother dyad | 1.96 | 0.10 to 5.43 | |
| Mother | 2.32 | 0.25 to 5.35 | |
| Litter | 1.55 | 0.07 to 4.31 | |
| | Estimate | 95% CI | Probability of direction (%) |
| B) Fifty meters | | | |
| Intercept | -5.89 | -11.41 to -3.40 | - |
| Maternal kinship 0.25 | 6.13 | 1.48 to 14.16 | 99.75 |
| Maternal kinship 0.5 | 6.31 | 1.35 to 14.19 | 99.30 |
| Genetic kinship | 0.91 | 0.04 to 2.19 | 98.02 |
| | SD | 95% CI | |
| Random effects | | | |
| Mother dyad | 1.96 | 0.10 to 5.43 | |
| Mother | 2.32 | 0.25 to 5.35 | |
| Litter | 1.55 | 0.07 to 4.31 | |

Table S4.

Output of the Bayesian Gamma mixed-effects model testing how den distance influences synchrony depending on kin status (N = 180 litter combinations with 85 litters from 54 mothers).

| | Estimate | 95%CI | Probability of direction (%) |
|---|---|---|---|
| Intercept | 4.02 | 3.86 to 4.18 | - |
| Maternal kinship: yes | -0.36 | -0.68 to -0.03 | 98.29 |
| | SD | 95%CI | |
| Smooth terms | | | |
| Den distance, maternal kinship = no | 0.31 | 0:01 to 1.53 | |
| Den distance, maternal kinship = yes | 1.23 | 0.06 to 4.05 | |
| Random effects | | | |
| Mother dyad | 0.11 | 0.00 to 0.33 | |
| Mother | 0.13 | 0.00 to 0.39 | |
| Litter | 0.12 | 0.01 to 0.35 | |

Table S5.

Model 2 - Output of the Bayesian Gamma mixed-effects model showing the effects of relationship strength and maternal kinship on the time between births within social groups, using the proximity measures (time observed within two body lengths) and CSI of affiliative behaviours (excluding proximity) from focal observations to approximate relationship strength (N = 35 litter combinations with 31 litters from 23 females). Proximity (mean = 3.6, sd = 4.3) and CSI (mean = 1.0, sd = 1.6) were z-transformed.

| | Estimate | | 95%CI | | |
|---|---|---|---|---|---|
| | Log scale | % change | Log scale | % change | Probability of direction (%) |
| Intercept | 4.05 | - | 3.19 to 4.73 | - | - |
| Proximity | -0.04 | -3.9 | -0.46 to 0.38 | -36.9 to 46.2 | 57.06 |
| CSI | -0.16 | -14.8 | -0.58 to 0.26 | -44.0 to 29.7 | 78.60 |
| Kinship 0.5 | -0.75 | -52.8 | -1.42 to -0.06 | -77.8 to -5.8 | 97.78 |
| Kinship 0.25 | -0.47 | -37.5 | -1.22 to 0.31 | -70.5 to 36.3 | 87.55 |
| | SD | | 95%CI | | |
| Smooth term | | | | | |
| Seasonality | 0.20 | | 0.01 to 0.63 | | |
| Random effects | | | | | |
| Mother dyad | 0.35 | | 0.01 to 1.05 | | |
| Mother | 0.62 | | 0.02 to 1.97 | | |
| Litter | 0.68 | | 0.02 to 2.03 | | |

Table S6.

Models 3 and 4 - Output of the Bayesian Bernoulli mixed-effects models analysing the effect of synchrony (i.e. days between births) on the occurrence and frequency of alloparental care. The outcome variables, i.e. respectively the presence of allonursing or the presence of other forms of allomaternal care (allogrooming and lying in body contact) were based on (**A**) binary data per dyad, collected during both focal and scan observations (N = 33 litter combinations with 42 litters from 32 females), and (**B**) binary data per focal session (N = 23 litter combinations with 22 litters from 17 females). Results are shown in log-odds. Days between births were z-transformed (A: mean = 46.1, sd = 36.1; B: mean = 45.0, sd = 37.0).

| | A) Occurrence of allomaternal care per dyad (N = 33) | | | | | |
|---|---|---|---|---|---|---|
| | Allonursing | | | Grooming and body contact | | |
| | Estimate | 95% CI | Probability of direction (%) | Estimate | 95% CI | Probability of direction (%) |
| Intercept | -2.14 | -4.78 to 0.39 | - | -0.76 | -3.53 to 1.88 | - |
| Days between births | -3.41 | -5.91 to -1.36 | 100 | -1.57 | -3.94 to 0.20 | 95.8 |
| Kinship 0.5 | 2.80 | -0.01 to 5.69 | 97.5 | 2.87 | -0.06 to 5.92 | 97.3 |
| Kinship 0.25 | 2.22 | -0.95 to 5.36 | 91.9 | 1.65 | -1.57 to 4.97 | 84.8 |
| Source scan | -2.89 | -5.91 to -0.07 | 97.7 | -3.26 | -6.15 to -0.46 | 98.8 |
| | SD | 95% CI | | SD | 95% CI | |
| Random effects | | | | | | |
| Mother dyad | 1.41 | 0.06 to 4.08 | | 2.25 | 0.11 to 6.28 | |
| Mother | 1.80 | 0.07 to 5.49 | | 2.54 | 0.10 to 7.92 | |
| Litter | 2.33 | 0.11 to 6.72 | | 2.43 | 0.08 to 8.40 | |
| | B) Occurrence of allomaternal care per focal session (N = 23) | | | | | |
| | Allonursing | | | Grooming and body contact | | |
| Intercept | -7.82 | -10.28 to -5.32 | - | -5.83 | -8.11 to -3.43 | - |
| Days between births | -1.79 | -3.63 to -0.23 | 99.0 | -0.33 | -1.79 to 0.95 | 68.9 |
| Kinship 0.5 | 2.09 | -0.35 to 4.64 | 95.4 | 1.89 | -0.51 to 4.47 | 94.0 |
| Kinship 0.25 | 1.67 | -1.12 to 4.49 | 88.5 | 0.84 | -1.98 to 3.76 | 72.1 |
| | SD | 95% CI | | SD | 95% CI | |
| Random effects | | | | | | |
| Mother dyad | 0.85 | 0.03 to 2.63 | | 1.24 | 0.06 to 3.26 | |
| Mother | 1.56 | 0.05 to 5.48 | | 2.48 | 0.14 to 6.66 | |
| Litter | 1.12 | 0.05 to 5.48 | | 1.69 | 0.40 to 4.17 | |

**Dataset S1 (separate file).** Data for the analysis of birth synchrony, using proximity data recorded during the scan observations to calculate relationship strength.

**Dataset S2 (separate file).** Data for the analysis of birth synchrony, using data on affiliative behaviours recorded during focal observations to calculate relationship strength.

**Dataset S3 (separate file).** Data for the analysis of the occurrence of allomaternal care, using ad lib data collected during both scan and focal observations.

**Dataset S4 (separate file).** Data for the analysis of the frequency of allomaternal care, using data collected during the focal observations.

**Dataset S5 (separate file).** Data for the analysis of the effect of oestrus synchrony on male monopolization.

**Script S1 (separate file).** Script for the analysis of birth synchrony, using proximity data recorded during the scan observations to calculate relationship strength.

**Script S2 (separate file).** Script for the analysis of birth synchrony, using data on affiliative behaviours recorded during focal observations to calculate relationship strength.

**Script S3 (separate file).** Script for the analysis of the occurrence of allomaternal care, using ad lib data collected during both scan and focal observations.

**Script S4 (separate file).** Script for the analysis of the frequency of allomaternal care, using data collected during the focal observations.

**Script S5 (separate file).** Script for the analysis of the effect of oestrus synchrony on male monopolization.

### SI References

1. Concannon PW. 2011 Reproductive cycles of the domestic bitch. *Animal Reproduction Science* **124**, 200–210. (doi:10.1016/j.anireprosci.2010.08.028)

2. Car C *et al.* 2025 Mating system of free-ranging domestic dogs and its consequences for dog evolution. *Proc. Natl. Acad. Sci. U.S.A.* **122**, e2421756122. (doi:10.1073/pnas.2421756122)

3. Belisle P, Chapais B. 2001 Tolerated co-feeding in relation to degree of kinship in Japanese macaques. *Behaviour* **138**, 487–509.

4. Silk JB. 2002 Kin selection in primate groups. *Int J Primatol* **23**, 849–875. (doi:10.1023/A:1015581016205)

5. Kappeler PM, van Schaik CP, editors. 2007 *Cooperation in primates and humans: Mechanisms and evolution*. Heidelberg: Springer-Verlag Berlin Heidelberg.

6. Smith JE. 2014 Hamilton's legacy: Kinship, cooperation and social tolerance in mammalian groups. *Animal Behaviour* **92**, 291–304. (doi:10.1016/j.anbehav.2014.02.029)

7. Chang CC, Chow CC, Tellier LC, Vattikuti S, Purcell SM, Lee JJ. 2015 Second-generation PLINK: Rising to the challenge of larger and richer datasets. *Gigascience* **4**, 7. (doi:10.1186/s13742-015-0047-8)

8. Berghänel A, Lazzaroni M, Ferenc M, Pilot M, El Berbri I, Marshall-Pescini S, Range F. 2025 Cofeeding at rich clumped food patches in free-ranging dogs: social tolerance or scramble competition? *Behav Ecol Sociobiol* **79**, 51. (doi:10.1007/s00265-025-03590-8)

9. Silk JB, Alberts SC, Altmann J. 2003 Social bonds of female baboons enhance infant survival. *Science* **302**, 1231–1234. (doi:10.1126/science.1088580)

10. Csárdi G, Nepusz T, Traag V, Horvát S, Zanini F, Noom D, Müller K, Schoch D, Salmon M. 2025 igraph: Network analysis and visualization in R. (doi:10.5281/zenodo.7682609)

11. Hoppitt WJE, Farine DR. 2018 Association indices for quantifying social relationships: How to deal with missing observations of individuals or groups. *Animal Behaviour* **136**, 227–238. (doi:10.1016/j.anbehav.2017.08.029)
